# The inevitability and superfluousness of cell types in spatial cognition

**DOI:** 10.1101/2024.01.10.575026

**Authors:** Xiaoliang Luo, Robert M. Mok, Bradley C. Love

## Abstract

Discoveries of functional cell types, exemplified by the cataloging of spatial cells in the hippocampal formation, are heralded as scientific breakthroughs. We question whether the identification of cell types based on human intuitions has scientific merit and suggest that “spatial cells” may arise in non-spatial computations of sufficient complexity. We show that deep neural networks (DNNs) for object recognition, which lack spatial grounding, contain numerous units resembling place, border, and head-direction cells. Strikingly, even untrained DNNs with randomized weights contained such units and support decoding of spatial information. Moreover, when these “spatial” units are excluded, spatial information can be decoded from the remaining DNN units, which highlights the superfluousness of cell types to spatial cognition. Now that large-scale simulations are feasible, the complexity of the brain should be respected and intuitive notions of cell type, which can be misleading and arise in any complex network, should be relegated to history.

## 1 Introduction

Spatial cognition encompasses our cognitive abilities to know where we are and navigate in complex environments, which includes self-localization, comprehension of environmental layouts, and efficient navigation toward distant goal locations. The prevailing view of the field attributes these functions to neural machinery within the hippocampal formation which contain a diverse array of “spatial” cell types including place cells, head-direction cells, border cells, and grid cells (e.g., Hartley et al. 2014; Moser et al. 2008; Grieves and Jeffery 2017), and that these cells collectively form the foundation of an internal cognitive map of space that underlie our spatial abilities (O’Keefe and Dostrovsky, 1971; Taube et al., 1990; Lever et al., 2009; Hafting et al., 2005). As such, a substantial proportion of the field focuses on characterizing particular functional “cell types” based on neural activity patterns in these regions. However, these “spatial cells” were identified due to their intriguing firing patterns that piqued the interest of neuroscientists and later defined based on subjective criteria – place cells appear to encode particular locations, border cells respond to borders, head-direction cells are sensitive to a heading direction, grid cells exhibit firing fields in a grid-like pattern. In practice, the field’s initial fascination with these cells has led to a stubborn tendency to focus on “cell type” classification and overlook the fact that the criteria for distinct cell types seldom align seamlessly with empirical data and often exhibit mixed selectivity across task and environmental variables (e.g., Hollup et al. 2001; McKenzie et al. 2013; Dupret et al. 2010; Eichenbaum 2015; Grieves and Jeffery 2017; Latuske et al. 2015; Tang et al. 2015), raising questions about their precise roles, if any.

Could these cells simply arise from any systems with complex computations, unrelated to processes specific to spatial or navigation? There are strong hints that “spatial” cells may arise simply from domain-general learning algorithms that can lead to both concept and spatial neural representations (Mok and Love, 2019, 2023; Stachenfeld et al., 2017; Whittington et al., 2020). Indeed, models show that spatial firing patterns can arise from factors entirely unrelated to spatial cognition, such as sparseness constraints (Kropff and Treves, 2008; Franzius et al., 2007a,b).

Another question is whether these cells are particularly important for spatial cognition. Research has shown that “spatial” cells may not be functionally more critical than those with other cells with multiplexed or hard-to-interpret firing patterns for spatial localization (Diehl et al., 2017) or for explaining neuronal response profiles in the medial entorhinal cortex (Nayebi et al., 2021). Even lesion studies in the hippocampal formation do not consistently result in specific navigation impairments (Hales et al., 2014; Whishaw and Tomie, 1997; Whishaw et al., 1997).

Considering that both biological and artificial “cells” rarely perfectly align with human-defined criteria with many cells exhibit “uninterpretable” firing patterns, one unsettling possibility is that the neuroscience may have inadvertently constrained its pursuit by fixating on the search for interpretable “cell types”. By adopting this top-down, perhaps näıve approach, the field may be investing a large proportion of its resources in a potentially fruitless quest, ignoring other explanations for the emergence of spatial cells and the foundations of spatial cognition.

A harsh appraisal of the field is that it looks for whatever cell type superficially reflects the answer sought. For example, how do animals localize themselves? The naive answer is that there must be place cells. Setting aside this answer provides zero insight into how such a cell came to be, there is no reason for the brain to perfectly align with our intuitions and, equally problematic, cells with these properties might arise but not serve the top-down function neuroscientist ascribe to them. Ascribing interpretable functions to cells and naming them may be good for neuroscience careers, but is it good for neuroscience?

In this contribution, we offer an alternative to the prevailing notion that cells with readily interpretable receptive fields in regions like hippocampus underlie spatial systems. We propose that what we commonly refer to as “spatial cells” might be inevitable derivatives of general computational mechanisms, and are in no way intrinsically spatial. Our results demonstrate that these cells could manifest within any sufficiently rich information processing system, such as perception. Remarkably, not only do we find spatial signals within a non-spatial system but also demonstrate that these purported signals, ostensibly representing spatial knowledge, do not appear to have a privileged role for tangible downstream tasks related to spatial cognition as previously claimed.

To evaluate these possibilities, we turned to deep neural networks (DNNs) that were information processing systems devoid of components dedicated to spatial processing. Specifically, we analyzed deep learning models of perception with hundreds of millions of parameters. DNNs trained on natural images have achieved human-level performance in recognizing real-world objects in photographs (Krizhevsky et al., 2012; Simonyan and Zisserman, 2015). These models exhibit strong parallels with representations in the primate ventral visual stream (Khaligh-Razavi and Kriegeskorte, 2014; Yamins et al., 2014; Güçlü and van Gerven, 2015; Eickenberg et al., 2017; Zeman et al., 2020). Our investigation aims to determine whether processing egocentric visual information within such complex – yet non-spatial – models can lead to brain-like representations of allocentric spatial environments, and whether these representations are essential for spatial cognition. Additionally, we considered whether untrained DNNs (absent any experiences) can account for the emergence of spatial cells and spatial cognition.

To evaluate functional spatial information in perceptual DNN models, we began by creating a three-dimensional virtual environment reminiscent of a laboratory used in animal studies in neuroscience (Fig. 1). In this virtual setting, an agent randomly forages in a two-dimensional square area, much like an animal would explore an enclosure. The agent “sees” images of the room within its field of view over many locations and heading directions. Images from the first-person perspective are processed by a DNN, and we assess whether the internal representations of the model contain various kinds of spatial knowledge. We adopted a decoding framework which parallels experimental techniques employed by neuroscientists. Specifically, we trained linear regression models using various levels of representations (unit activations from visual inputs) generated by DNNs of different architectures to decode navigation-relevant variables, including self-localization, heading direction, and distance to the closest wall on locations and heading directions they have not encountered before. To test whether DNN units carry information like “spatial” cells in the brain, we classified and visualized model units based on standard criteria for place, direction, and border cells (Tanni et al., 2022; Banino et al., 2018). To evaluate whether these cells play a privileged role in spatial cognition, we excluded units classified as spatial units based on traditional spatial cell criteria and assessed their spatial knowledge.

**Figure 1:**
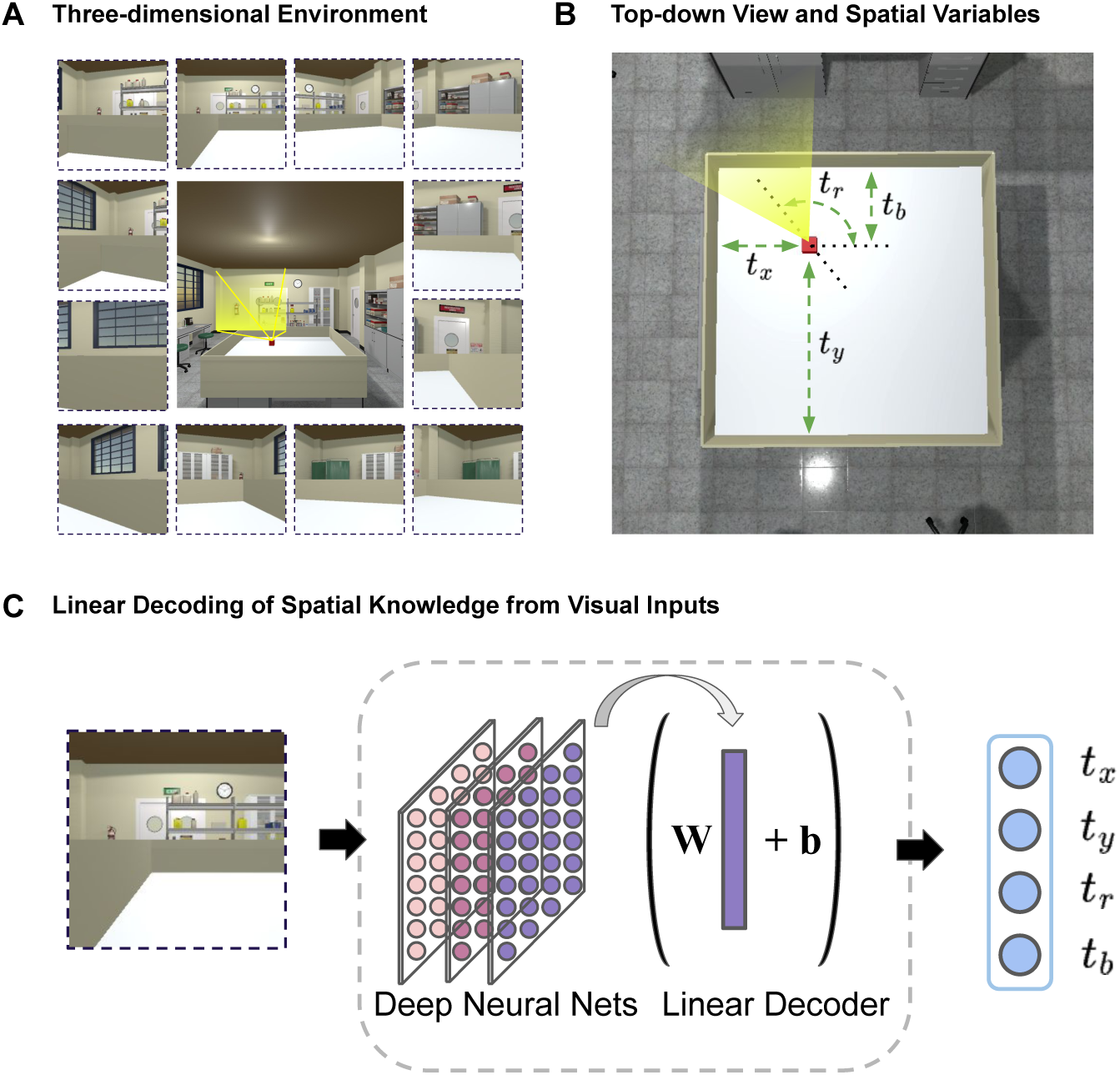
Assessing spatial knowledge in non-spatial perception systems using a linear decoding approach in a virtual environment. (A) A three-dimensional virtual space is created to resemble a realistic laboratory environment with a variety of visual features. An agent moves randomly in a two-dimensional area which is within a three-dimensional space and processes first-person views of the environment. Central image shows the threedimensional environment. Surrounding images are example views taken by the agent at different locations and heading directions. (B) Top-down view of the area where the agent can explore. We define spatial knowledge of the agent with four values. The agent’s location is denoted by the Cartesian coordinates t*_x_*, t*_y_*. The agent’s heading direction is denoted by the angle t*_r_*. The distance between the agent and the nearest wall is t*_b_*. (C) Individual views are processed by perception models (deep neural networks of object recognition). We train linear regression models with various levels of internal representations from these networks to assess spatial knowledge related to self-location (t*_x_*, t*_y_*), heading direction (t*_r_*) and distance to the closest wall (t*_b_*).

To foreshadow our findings, we discovered that DNNs, regardless of their architectural variations, representation levels, or training states, possess noteworthy proficiency in allocentric spatial understanding. Spatial variables including location, heading direction and distance to boundaries can be accurately decoded from the DNN models. Additionally, a substantial number of DNN units exhibit spatial firing patterns that align with the criteria used in neuroscience to classify place, head-direction, and border cells. Notably, a significant portion of these units employs mixed-selectivity and conjunctive coding, akin to characteristics found in hippocampal neurons (Eichenbaum, 2015). We further revealed that excluding units meeting the classical criteria of spatial cell types has minimal impact on spatial decoding performance, reinforcing the perspective that the code underpinning spatial cognition is more distributed and diverse than previously thought, relying on neural population coding rather than individual and specialized cell types (Hebb, 1949; Eichenbaum, 2018; Ebitz and Hayden, 2021). In summary, our analyses suggest that “spatial” cell types may be inevitable and superfluous in complex information processing systems.

## 2 Results

### 2.1 Spatial Knowledge through Non-spatial Systems

We predict that spatial knowledge can arise in computational systems with sufficient complexity absent a spatial grounding. We test this hypothesis by analyzing the percepts of an agent freely moving in a virtual environment. The percepts take the form of viewpoints fed into a deep neural network with its millions of weights either randomly initialized or trained for object recognition, a non-spatial task. From various network layers, we attempted to decode spatial information such as location, from this complex computational system.

To assess information latent in complex networks, we trained linear regression models to decode the agent’s location, head-direction and distance to the closet border using fixed unit activity extracted at various levels of the networks. While sequential information is important for spatial systems, our decoding framework does not utilize sequential information of sampled views during training, making it a stricter test for the central claim that spatial knowledge can form out of general information processing. We expect that one’s location, heading-direction and distance from border may be decodable from visual information alone due to the inherent continuity of perception across spatial locations and/or directions. All results reported here are tested on out-of-sample locations and views using the perception model VGG-16 (Simonyan et al., 2014), unless noted otherwise. For details referring to training and testing procedure, see Methods. For results of other models such as Resnet-50 (He et al., 2016) and Vision Transformers (ViT; Dosovitskiy et al. 2020), see Appendix.

We find that spatial knowledge, namely one’s own location (spatial coordinates; Fig. 2B, left panel), heading direction (Fig. 2B, middle panel) and distance to the nearest wall (Fig. 2B, right panel) can be successfully decoded from unit activity in a perception model. We quantified the degree of spatial knowledge by the decoding error, that is, the deviation of the model’s prediction of its own location, heading direction, or distance to the nearest wall relative to the ground truth (i.e., the physical unit of space or angle in the virtual environment). We normalized decoding error with respect to the maximal unit length of the moving area (see Methods). For example, a normalized error of 0.05 for location prediction means the decoded location is 5% off from the true location relative to the maximal width/depth of the entire moving area. As more locations and views are sampled for training the agent (x-axis), decoding performance improves. The agent demonstrates impressive decoding performance across all spatial knowledge tests, even when trained on just 30% of randomly selected independent locations and associated viewpoints within the entire space. The moving area encompasses 289 distinct locations, each featuring 24 evenly spaced views covering the full 360-degree spectrum (for more details, please see Methods).

**Figure 2:**
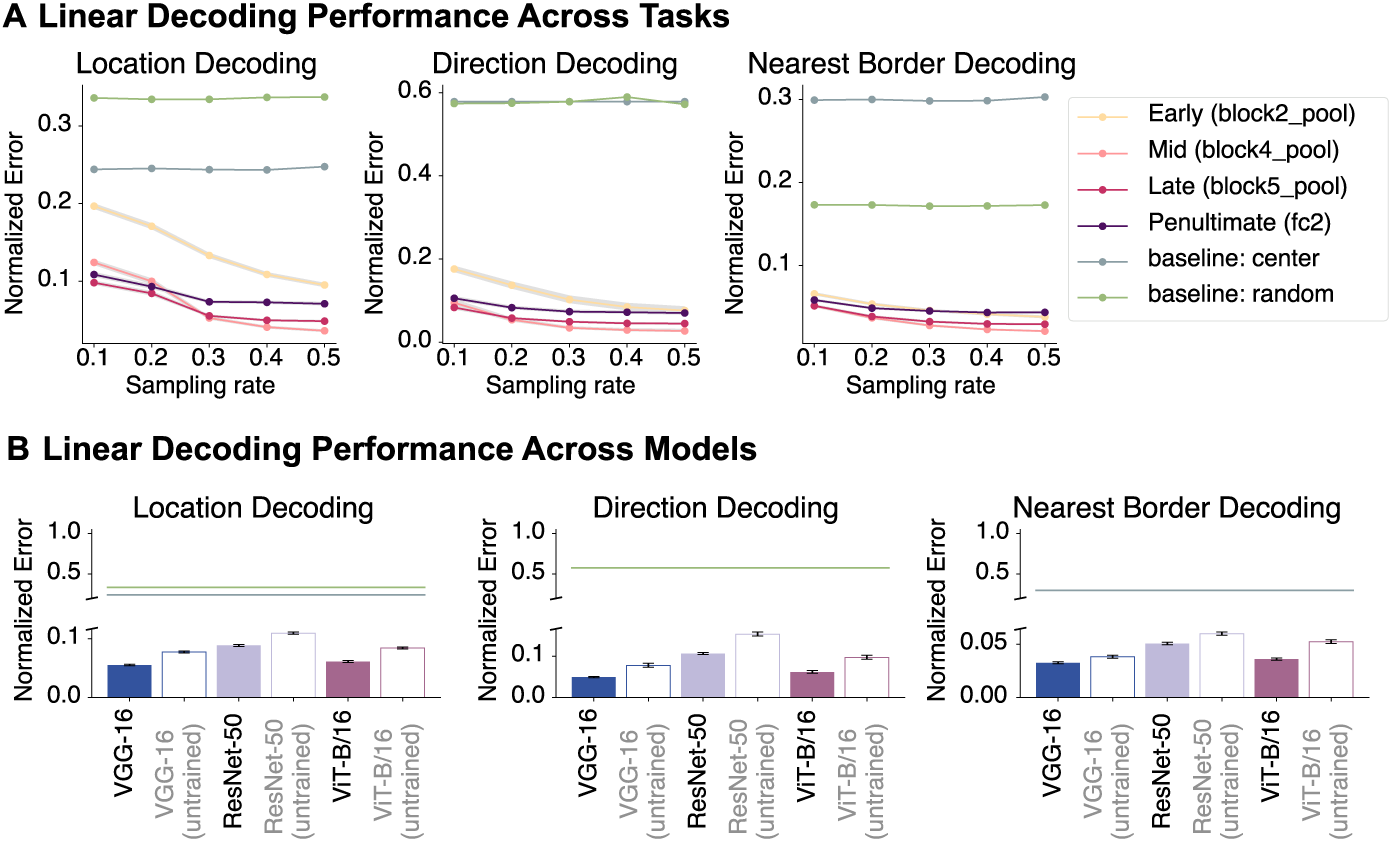
Perception models absent a spatial basis possess extensive spatial knowledge. (A) Decoding performance across tasks and model layers. Across three tasks, mid-to-late layers exhibited lower decoding errors compared to early layers and the penultimate layer of VGG-16 (fc2). All layers outperformed chance (green and blue lines). (B) Decoding performance across a number of deep neural network architectures (penultimate layer; see Appendix for full results), including convolutional networks and vision transformers. All pre-trained and untrained models outperformed baseline measures. Error is in normalized virtual environment units (see Methods). See full results across models, layers and sampling rates in the Appendix. Shaded areas in B and error bars in C represent 95% confidence intervals of the mean decoding error (bootstrapped across locations).

Decoding performance varied across layers of the network where deeper layers typically achieved lower decoding error (apart from the penultimate layer in VGG-16, i.e., fc2). Crucially, decoders across all sampling rates and levels of representation show substantially better performance than chance (green and blue lines), determined by strategies invariant to visual inputs (i.e., random or predicting the center of possible choices; see Methods). Our results demonstrate there is sufficient spatial information to achieve remarkably low decoding error in a perception model based on a few visual snapshots of the virtual space, which could be further improved through the integration of information across views in a perception-based navigation system.

To test the generality our claim that spatial knowledge can arise from complex computational systems irrespective of their architectural variations or training states, we examined several computational systems including deep convolutional neural networks (DCNN) and Vision Transformers (ViT). We evaluated these models in both their pre-trained form (optimized for visual object recognition), and in an untrained state (randomly initialized parameters), following the same decoding approach described earlier. Here, we present the results for the penultimate layer representation and uniformly sampled 30% of all locations and their views for training the decoder, but similar results were observed across layers (for full results across various models, layers, and sampling rates, see Appendix.). We found that all models possessed latent spatial knowledge, exhibiting lower errors far below chance (Figure 2C). Notably, this was true even in untrained networks. While one might contend that deep convolutional neural networks, with their reliance on local convolution operations, inherently possess a predisposition for extracting spatial knowledge from visual information, the revelation that an untrained ViT—comprising nothing more than a hierarchy of fully-connected layers (i.e., self-attention) and non-linear operations—can achieve superior decoding performance illustrates the inevitability of computational complexity in facilitating the extraction of spatial information necessary for cognitive spatial processes, even in non-spatial systems.

### 2.2 “Spatial Cells” Inevitably Arise in Complex Computational Systems

How was it possible to decode spatial information from non-spatial networks? The field’s common conception is that cells exhibiting spatial firing profiles play a pivotal role in supporting spatial cognition, as these profiles appear intuitively useful to spatial tasks such as navigation. Could spatial cells be responsible for the impressive decoding of spatial information within a perception system like we have shown? If a perception system devoid of a spatial component demonstrates classically spatially-tuned unit representations, such as place, head-direction, and border cells, can “spatial cells” truly be regarded as “spatial”? Might they be inevitable derivatives of any complex system? If so, do they drive spatial knowledge, or are they in fact superfluous in spatial cognition?

To determine if the model’s spatial knowledge is primarily supported by spatially-tuned units like those found in the brain, we classified every hidden unit in each model based on criteria used to identify spatial cells in neuroscience for place cells (Tanni et al., 2022), head-direction cells (Banino et al., 2018), and border cells (Banino et al., 2018), respectively (see Methods). The composition of cell types across layers is shown in Fig. 3A. Contrary to the predominant view that “spatial cells” are unique properties of spatial systems and are grounded in space-related tasks, we found many units in the non-spatial model VGG-16 that satisfied the criteria of place, head-direction, and border cells (see Methods for criteria; for other models, see Appendix). The majority of units show mixed selectivity irrespective of layer depth. Notably, a significant proportion of units show both place and directional tuning (P+D), matching a common observation in the literature (e.g., McNaughton et al. 1991; Tang et al. 2015; Grieves and Jeffery 2017). We selected example units and plotted their spatial activation patterns in the two-dimensional virtual space (irrespective of direction), and their direction selectivity in polar plots (Fig. 3B-3E; for more examples, see Appendix). These examples show spatially-tuned units without directional tuning that match hippocampal place cells (Fig. 3B), direction-selective units with strong direction selectivity accompanied by minor spatial selectivity matching head-directional cells (Fig. 3C), and units with boundary cell-like tuning (Fig. 3D). There were also many units that exhibited both strong place and directional tuning (Fig. 3E). These results support our theory that “spatial cells” might arise in any computational systems, even in systems designed for nonspatial tasks (e.g., object recognition). Consistent with our view, we found no clear relationship between cell type distribution and spatial information in each layer. This raises the possibility that “spatial cells” do not play a pivotal role in spatial tasks as is broadly assumed.

**Figure 3:**
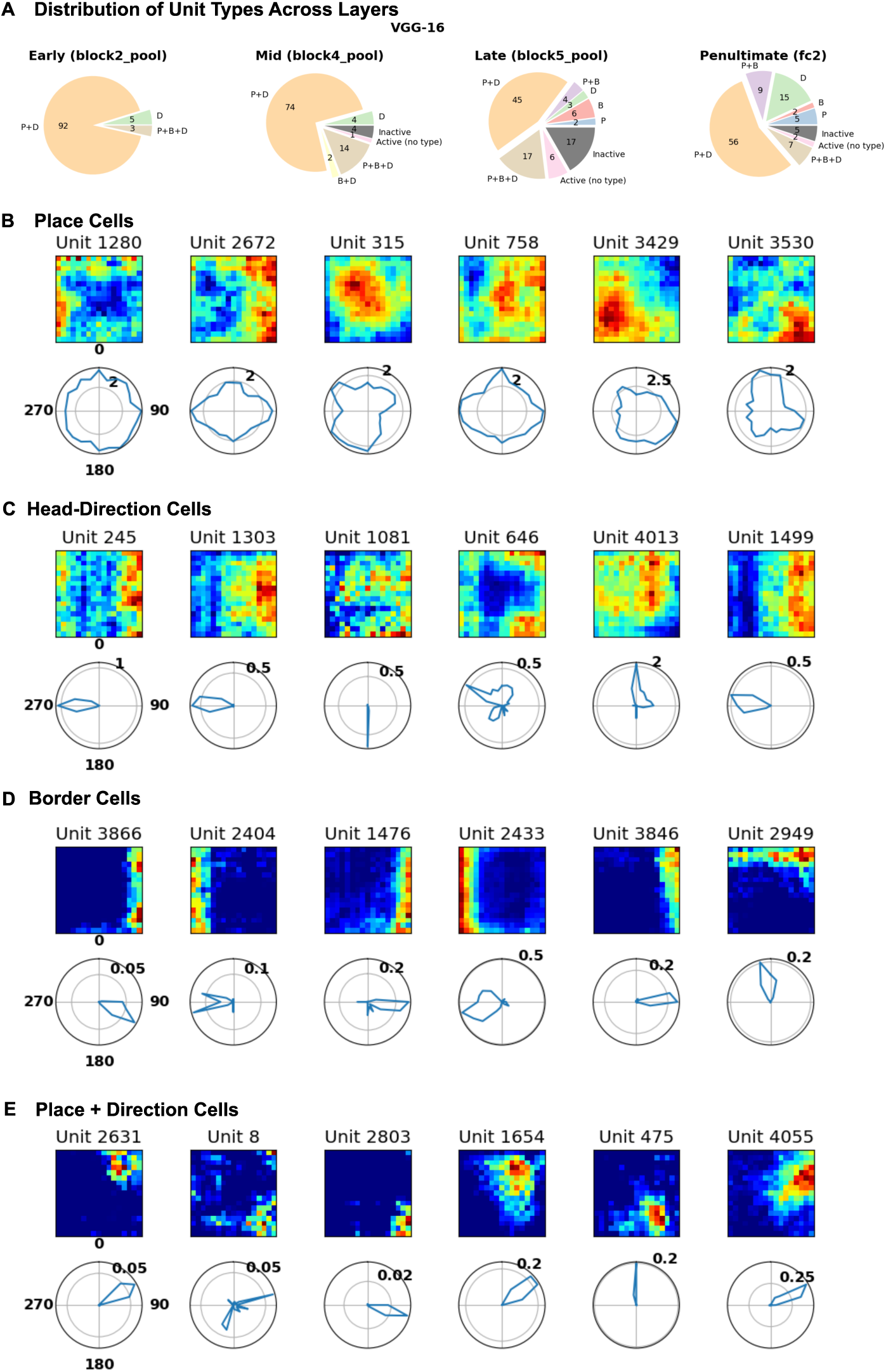
Representations of perception models developed for object recognition exhibit typical spatial cell-like firing profiles. Active units were classified across model layers based on standard criteria used to identify place cells (P), head-direction cells (D), and border cells (B). Example units from VGG-16 (see Appendix for more examples from other models). (A) Pie charts illustrating the proportion of “spatial” cell types identified in DNNs, including units that are inactive. Many units satisfied the criteria for place, head-direction and border cells irrespective of layer depth. Many units exhibited mixed selectivity, with a significant amount of units displaying strong place and directional tuning. (B-E) Spatial firing profiles of model units in spatial activation maps and polar plots . For activation maps, each unit’s activation was plotted at each location in the two-dimensional area irrespective of heading direction. For direction selectivity, polar plots show the average activity of the activity map across location at a given angle, which reflects the tuning magnitude to each heading direction). (B) Example place cell units that show strong spatial selecitivity with little direction selectivity. (C) Examples of head-direction cell units that show strong direction selectivity but weak location selectivity. (D) Examples of border cell units that respond strongly to boundaries of the environment. (E) Examples of mixed-selective place and head-direction cell units with strong spatial and directional tuning. Examples presented are from VGG-16. See Appendix for more examples.

### 2.3 “Spatial Cells” Are Superfluous in Spatial Cognition

We have establishd that“spatial cells” can arise in non-spatial systems. Here, we consider whether units with spatial properties are necessary to decode spatial information. It may very well be that spatial units, akin to place cells, might not only arise in complex (including random) networks but are also superfluous to spatial cognition. To test this possibility, we performed a systematic exclusion analysis to assess whether model units that exhibit the strongest spatial firing properties contribute to spatial knowledge. First, we scored and ranked each unit classified as place, head-direction, and border cell units (see Methods), and then re-trained linear decoders without the top n units of a specific cell type and evaluated the model’s corresponding spatial knowledge of the environment. That is, we test model’s location, heading direction and distance to closet border decoding ability by excluding place, head-direction and border cell units, respectively. We repeated this procedure with a progressively higher exclusion ratio (see Methods).

In line with our hypothesis, excluding spatial units in the models that scored highest on each of the corresponding criteria (place field activity, number of place fields, directional tuning, and border tuning), had minimal effect on decoding performance even with a large proportion of the highest ranked spatial units being excluded (Fig 4B, top panel). To assess whether any detriment was observed was significant and specific to excluding highly-ranked spatial units, we randomly excluded the same number of units without regard to the spatial score (Fig. 4B, bottom panel) and observed a similar pattern across four exclusion scenarios, meaning that highly-tuned spatial units contributed no more to spatial cognition than a random selection of units.

**Figure 4:**
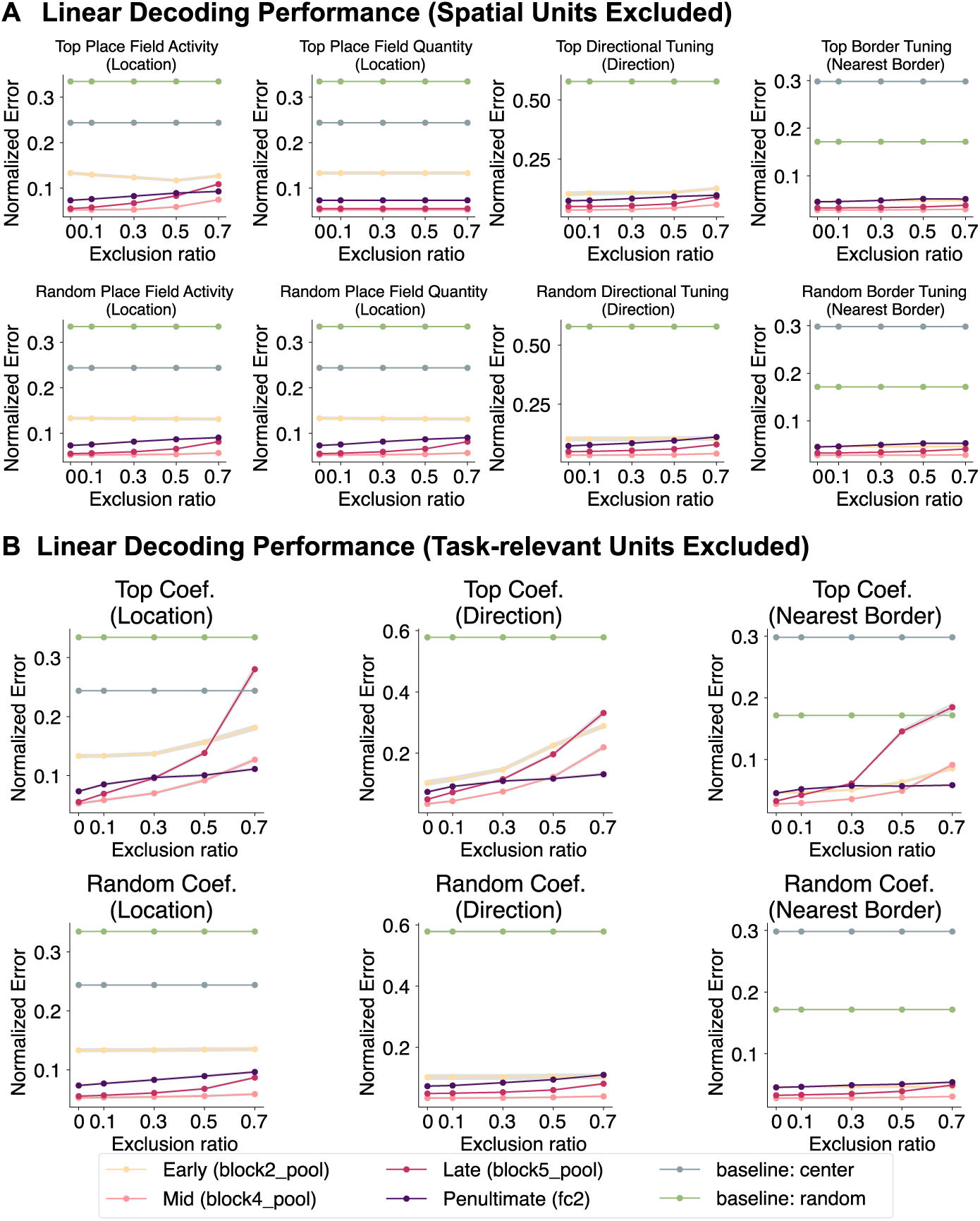
“Spatial” cells do not play a privileged role in spatial cognition. Exclusion analyses showed that units exhibiting traditional spatial firing profiles do not form the basis of the model’s spatial knowledge. (A) Units in each layer were ranked based on standard criteria for place (maximum place field activity, number of place fields), head-direction (strength of directional tuning), and border (strength of border tuning) cells. As more highly-ranked spatial units were excluded, the overall decoding performance remained relatively stable across all tasks (top). Excluding an equivalent number of units randomly yielded similar performance (bottom). (B) Model units’ contribution to spatial knowledge based on their contribution to decoding performance (magnitude of regression coefficients). More highly-ranked task-relevant units excluded results in a marked deterioration of decoding performance by decoders trained on the remaining units (top). Randomly excluding an equivalent number of units had minimal impact on performance (bottom). Results here are from VGG-16. For other models, please refer to the Appendix.

Finally, we performed an additional analysis where we excluded model units that contributed most to each of the tasks. Specifically, we identified each unit’s importance for each task, based on the magnitudes of the coefficients learned by the linear decoders (trained in Section 2.1), excluded the top n units, and re-trained linear decoders for each respective task using the remaining units. This approach had a greater impact on decoding error compared to spatial unit criteria or random exclusion, with performance suffering more as the exclusion ratio rose (see Figure 4C). Nevertheless, decoding was still possible at high exclusion ratios, suggesting that spatial information is widely distributed across network units.

## 3 Discussion

The prevailing perspective in neuroscience conflates spatial knowledge and the variety of spatial cognitive abilities in animals with “spatial cells” in the hippocampal formation. In this work, we proposed an alternative perspective where “spatial cells” can simply arise in non-spatial computational systems with sufficient complexity, and that these units may, in fact, be by-products of such systems absent any leading role in spatial cognition.

Deep neural networks (DNNs) for object recognition served as an ideal testbed for our hypotheses. Object recognition DNNs, designed to generate translation-invariant visual features for successful categorization, lack inherent spatial elements. Unlike spatial navigation systems (e.g., Banino et al. 2018; Cueva and Wei 2018), DNNs in object recognition do not purposefully incorporate information about velocity, heading direction, or event sequences. Exploring various DNN architectures for object recognition allows a comprehensive examination of the connection between spatial representations and non-spatial complex systems.

We applied DNNs to a three-dimensional virtual environment to simulate an agent foraging in a two-dimensional space, akin to how an animal explores an enclosure. Detailed analyses revealed that DNN’s representations contained sufficient spatial information to decode key variables, despite these DNNs being non-spatial perceptual systems. We found that many DNN units passed established criteria for classification as place, head-direction, and border cells, suggesting such units can be easily found in various non-spatial computational systems. Furthermore, excluding the “spatial” units had minimal impact on spatial decoding tasks, suggesting a non-essential role for spatial cognition. These results call into question utility of labelling these cell “types”.

We considered object recognition models from two prominent DNN families: deep convolutional networks, drawing inspiration from the mammalian visual system (Fukushima, 1980), and visual transformers, adapted from the Transformer architecture originally designed for natural language processing (Vaswani et al., 2017). Spatial information was readily decoded from models from both families, including untrained models with randomly initialized weights. The decoding results from the untrained visual transformer are particularly striking because transformers incorporate minimal inductive biases, such as constraints that respect spatial proximity (e.g., convolutions). In summary, complex networks that are not spatial systems, coupled with environmental input, appear sufficient to decode spatial information.

While previous modeling efforts have highlighted the importance of environmental sensory input in driving the emergence of spatial cells (e.g., Franzius et al. 2007a), it is worth noting the substantial theoretical and modeling differences to our contributions despite perceived similarity. Franzius et al. (2007) propose a sensory-driven model where spatial cells, assumed to be inherently special, emerge from visual input statistics tied to motion and slow-feature analysis, yet their model does not simultaneously produce place and head-direction cells nor assess their functional utility. In contrast, our work challenges the specialness of spatial cells, demonstrating their emergence in a non-spatial, nonsequential null model with fixed weights, while showing that these cells, though present, offer little benefit for spatial decoding tasks.

Indeed, our work raises the question of whether focusing on single cells and their “type” is the correct approach to understanding spatial cognition, or indeed any cognitive function. The field has maintained a persistent inclination towards identifying cell types based on firing profiles that are subjectively interpreted to be useful for the behavior of interest (Hartley et al., 2014; Moser et al., 2008; Sarel et al., 2017; Tsao et al., 2018; Høydal et al., 2019; Grieves et al., 2020; Ormond and O’Keefe, 2022). This is likely due to the historical backdrop in neuroscientific research where pioneers in visual neuroscience discovered and named individual cells in V1 by observing their firing profiles (Hubel and Wiesel, 1959) which won the Nobel Prize (1981) and marked a foundational era for the field. This paradigm, led to discoveries of place and grid cells (Nobel Prize in Physiology or Medicine, 2014), and the “Jennifer Aniston” neuron or “concept cells” (Quiroga et al., 2005), which were intuitively compelling and attractive as an apparent explanation. These cells may in fact play a role, but our work suggests that this must be critically assessed rather than assumed, as they may not play a privileged role compared to other cells in the population, and could even be superfluous for the context at hand. The general assumption that the simplicity and interpretability of firing profiles somehow lends support to a crucial role of these cells should be questioned. Unlike structurally or genetically defined cell types (e.g., pyramidal neurons, interneurons, dopamingeric neurons, c-Fos expressing neurons), it is unclear whether “cell types” in the spatial or even conceptual domain should be considered cell types in the same way. Indeed, the identification criteria for cell types themselves are often problematic and inconsistent across studies. For instance, place field selection thresholds vary from 20% to 25% of peak firing rate (Tanaka et al., 2018; Dombeck et al., 2010), speed thresholds range dramatically from 2 cm/s to 8.3 cm/s (Tanaka et al., 2018; Dombeck et al., 2010), and spatial bin requirements differ across leading studies (Tanni et al., 2022; Tanaka et al., 2018). As Grijseels et al. (2021) demonstrated, different detection methods produce vastly different place cell counts with minimal overlap between identified populations. This inconsistency raises critical questions about whether researchers are studying the same phenomena or entirely different neural populations.

With a growing recognition of the imperative to move beyond mere cell type categorization, as underscored by Olshausen and Field (2006) on the insufficient characterization of a majority of V1 cells, there is a new shift to focus on a broader perspective where neural assemblies or populations form the fundamental computational unit (Hebb, 1949; Eichenbaum, 2018; Ebitz and Hayden, 2021).

We pose a broader question: do our preconceptions of how complex systems *should* work hinder our aim to understand the brain’s inner workings? Notably, spatial representations conventionally linked to the hippocampal formation have been found in sensory areas (Saleem et al., 2018; Long et al., 2021; Long and Zhang, 2021). While prior attributions pointed to modulatory signals from the hippocampus, our study raises concerns about potential biases stemming from preconceived notions and an underestimation of the complexity inherent in the sensory system—it may be shouldering more extensive responsibilities than initially presumed. Work in other domains are reaching similar conclusions (cf. McMahon and Isik 2023).

It is also worth noting that the null model perspective extends beyond static representations to dynamic neural phenomena. Consider replay–the reactivation of neural firing patterns during rest or sleep, widely interpreted as evidence of memory consolidation and plasticity-dependent learning processes (e.g., Wilson and McNaughton 1994; Schapiro et al. 2018; Barry and Love 2022. However, our framework suggests that such sequential reactivation patterns could emerge as byproducts of complex system dynamics, independent of learning mechanisms. Just as spatial cell types manifest in untrained networks, replay-like patterns may arise from the inherent complexity of neural circuits without necessitating plasticity-driven explanations. Indeed, the key idea behind reservoir computing is that a pool of units with complex dynamics can provide the basis for useful computations in the absence of any plasticity (Jaeger and Haas, 2004). The mere observation of repeated activity patterns should not be conflated with evidence of learning-dependent processes.

One upshot of our contribution is that the pursuit of identifying and cataloging cell types aligned with subjective notions of how the brain works might impede scientific progress. While searching for cell types was a reasonable strategy in an era before it was straightforward to conduct large-scale simulations, this quest should now be relegated to history. Otherwise, apparent discoveries of cell types are likely to reflect the activity of broad classes of complex networks that implement a variety of functions or none at all in the case of the random networks we considered. It is far too easy for neuroscientists, including computational neuroscientists who design their models to manifest a variety of cell types, to fool themselves when the underlying complexity of the systems considered is not taking into account.

## 4 Methods

### 4.1 Virtual environment

We setup our virtual environment using the Unity3D Engine (editor version: 2021.3.16f1; Silicon). The three-dimensional laboratory environment was adapted from an existing template purchased on the Unity Store (3DEverything, 2022). Our virtual agent’s movement is constrained in a two-dimensional square placed inside the three-dimensional space, much like a moving area for an animal in real-world experiments. The agent randomly moves around the square area and captures first-person pictures of the environment across locations and heading directions.

### Environment specifications

The specifications for the environment, measured in Unity units, are as follows: the lab room has a height of 3.17 units, a width of 6.29 units, and a depth of 10.81 units. The moving area, which is a distinct part of the environment, has a height of 0.25 units, a width of 2 units, and a depth of 2 units. Additionally, the moving area is located at a specific location whose relative distances from the center to the lab room boundaries are: 3.59 units to the left wall, 5.35 units to the right wall, 5.55 units to the bottom wall, and 2.80 units to the front wall.

The agent is only allowed to move within the squared moving area. For simplicity, we discretize reachable locations by the agent in the area as a 2D grid where the agent can only move between points on the grid. We use a 17×17 grid (289 unique locations).

### Visual input

To collect visual inputs from the virtual environment, the agent randomly moves along the specified n-by-n grid in the squared area. We denote l as the total number of unique locations on the grid. The grid is evenly spaced. At each location, the agent turns around and captures m number of frames, each separated by a fixed angle (e.g., 15 degrees per frame). Each frame obtained by the agent is processed by a DNN (up to a given layer). Resulting outputs at the same location are considered independent data-points, each forming a long 1D vector. Repeating this procedure for all locations yields a M-by-F data matrix **X** where M = l × m and F is the number of feature variables. Each feature variable represents a neuron on the output layer of the DNN.

### 4.2 Spatial decoding

To evaluate whether representations in the deep neural networks (DNN) optimized for object recognition on natural images encode spatial information, we set up a series of spatial decoding tasks where we use fixed network representations across various hidden layers to decode spatial location (i.e., coordinates), head-direction (i.e., rotation degrees) and distance (i.e., Euclidean) to the nearest border using visual inputs of views from unvisited locations and/or directions. We formulate spatial decoding as a prediction problem where we fit a set of linear regression models on representations of visual views to predict location, direction and distance to nearest borders. We define the nearest border to the agent as the border that has the shortest Euclidean distance to the agent’s location regardless of the agent’s heading direction.

### Decoder training and testing

To fit and test spatial decoders, we randomly sample locations in the environment (subject to a sampling rate). When a location is sampled for training, we consider views of different rotations as independent data-points. For example, a location decoder is fit to predict a vector **t_i_** = [t*_x_*, t*_y_*, t*_r_*, t*_b_*] where t*_x_*, t*_y_* are the x, y coordinates, t*_r_* is the direction, t*_b_* is the shortest distance to a border, given a view respectively. For the entire training set (**X_i_**, **T_i_**) where **T_i_** = {**t_1_**, **t_2_**, …, **t_i_**}, we can write the decoder as

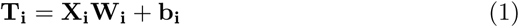

where **W_i_** are the coefficients and **b_i_** are the intercepts. Decoder weights (both coefficients and intercepts) are optimized to minimize the mean squared error between the true and predicted targets.

### Feature representation and selection

We consider a number of representations for visual inputs. Given a DNN model, we train separate decoders using representations across different layers. We apply L_2_ regularization on the decoder weights to reduce overfitting. Additionally, we consider different exclusion strategies (see Sec. 4.4) where unit representations are selectively removed based on their firing profiles (e.g., the number of place fields a unit contains).

### Performance measure

We report the normalized decoding error on the heldout set of location, head-direction and nearest border decodings for each representation at a given sampling rate. The normalized decoding error reflects the deviation of the model’s prediction on location, heading direction or distance to the nearest wall relative to the ground truth (i.e., in terms of physical unit of space or angle in the virtual environment) and normalized with respect to the maximal length of the moving area (2 units in both depth and width). To reflect variance of the average error, we compute a two-sided 95% confidence interval of the average using a bootstrapping approach over locations.

### Baselines

We compare decoding performance achieved by our agent to baselines where the agent decodes location, head-direction and distance to border based on fixed policies independent to visual inputs. Specifically, we establish two baselines where the agent simply either predict randomly within the legal bound or predict the centre position of the room (when predicting location), predict 90 degrees (when predicting rotation) and predict the distance from the centre of the room (when predicting nearest border distance).

### 4.3 Profiling spatial properties of model units

To gain a more intuitive understanding of the kinds of model units that are most useful for spatial decoding tasks, we profile all model units from the layers we consider based on their spatial patterns of firing. We consider the most common cell types that are found to encode spatial information as defined in the neuroscience literature, which include place cells, border cells and head-direction cells. For each cell type, we track a number of measures defined below.

### Place cells

In our investigation, the identification of place-cell-like model units hinges on several key characteristics of unit firing. Firstly, we examine the number of place fields exhibited by each unit, defining a qualified field as one that spans 2D space ranging from 10 pixels to half the size of the environment, as outlined in Tanni et al. (2022). Additionally, we consider the maximal activation of each field, providing insights into the intensity and significance of unit firing.

### Head-direction cells

Following Banino et al. (2018), the degree of directional tuning exhibited by each unit was assessed using the length of the resultant vector of the directional activity map. In our case, there is an activity map spanning entire 2D space for each direction. Vectors corresponding to each direction of an activity map were created:

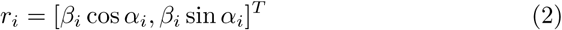

where α and β are, respectively, the angle and average intensity (irrespective of spatial locations) of direction i in the activity map. These vectors were averaged to generate a mean resultant vector:

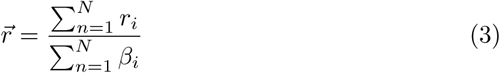

and the length of the resultant vector calculated as the magnitude of r⃗. We used 24 angular directions uniformly spaced.

### Border cells

Border score definition is based on Banino et al. (2018) where for each of the four walls in the square enclosure, the average activation for that wall, b*_i_*, was compared to the average centre activity c obtaining a border score for that wall, and the maximum was used as the border-score for the unit:

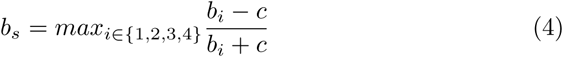

where b*_i_* is the mean activation for bins within d*_b_* distance from the i-th wall and c the average activity for bins further than d*_b_* bins from any wall (d*_b_* = 3). Units with border score > 0.5 are considered border-like.

### 4.4 Spatial decoding with unit exclusion

To delve deeper into how different units within the DNN models contribute to the linear decoding of spatial information, we employ two complementary analyses. Model units are selectively excluded subject to criteria detailed below. Spatial decoders are re-trained on the rest of the model units following the same procedure as before (sec. 4.2).

**Excluding units by spatial profile** In this first analysis, we selectively exclude units based on their firing profiles, which fall into the various spatial unit categories mentioned earlier. We initially rank all units in descending order according to their specific spatial measures (e.g., field count) and then systematically increase the exclusion rate, assessing the impact on the corresponding downstream tasks as we go. For example, we progressively exclude the top n% of units with the most place-like characteristics for varying values of n, examining the relationship between exclusion and performance of location decoding. This process is carried out separately for each unit type and their associated tasks we have defined, and we also include a control group where an equivalent number of units are excluded at random.

**Excluding units by task-relevance** Our second exclusion analysis focuses on identifying the task-relevance of each model unit by utilizing previously learned decoders and their coefficient magnitudes. Essentially, units with larger coefficients are deemed more crucial for decoding, while those with smaller or zero coefficients are considered less relevant. Similar to the exclusion by spatial profile, we gradually increase the exclusion rate by selectively removing units based on the magnitude of their decoder coefficients. Additionally, we conduct a control analysis where an equal number of units are randomly chosen for exclusion. We then retrain linear decoders with the remaining units and assess their performance on downstream spatial tasks.

## Data and code availability

The code for simulations and analyses are publicly available at https://github.com/dontpanic/Space.

## Acknowledgements

This work was supported by ESRC (ES/W007347/1), Wellcome Trust (WT106931MA), and a Royal Society Wolfson Fellowship (18302) to B.C.L., and the Medical Research Council UK (MC UU 00030/7) and a Leverhulme Trust Early Career Fellowship (Leverhulme Trust, Isaac Newton Trust: SUAI/053 G100773, SUAI/056 G105620, ECF-2019-110) to R.M.M.

## **A** Spatial decoding performance across various percetion models

**Figure S1:**
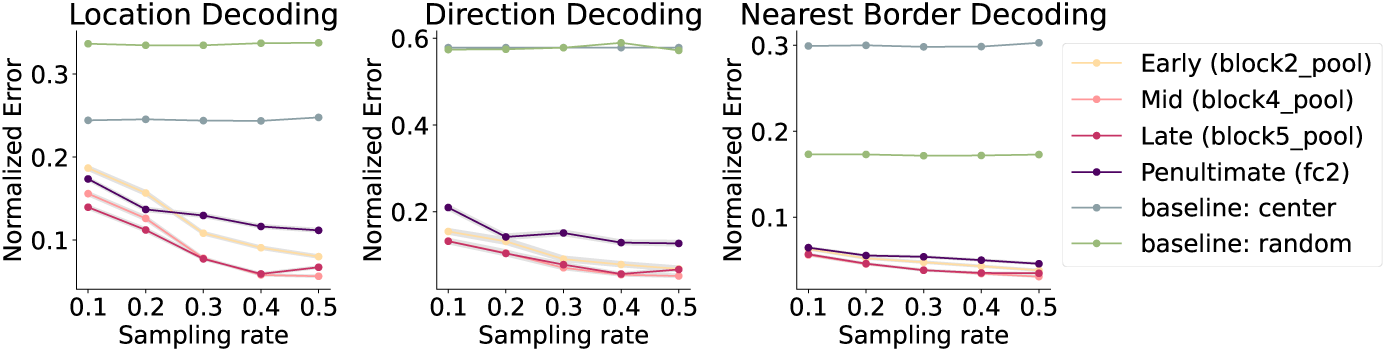
VGG-16 (untrained). Linear decoders trained with representations from various layers of the untrained VGG-16 achieve low errors across sampling rates; though not as good as its trained version. Mid-to-advanced layers show superior performance than early layers. As more locations are sampled for training the linear decoders, overall decoding performance improves. All model layers can decode better than two visual-invariant baselines.

**Figure S2:**
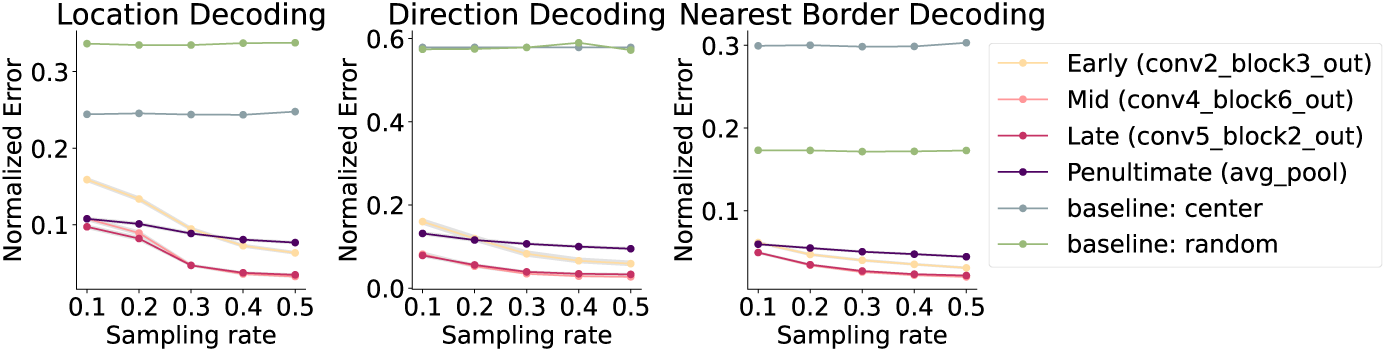
ResNet-50 (trained). Linear decoders trained with representations from various layers of the ResNet-50 pretrained on object recognition achieve low errors across sampling rates. Mid-to-advanced layers show superior performance than early layers. The penultimate layer decoding performance did not improve as much as the intermediate layers as more locations are sampled for training the linear decoders. All model layers can decode better than two visual-invariant baselines.

**Figure S3:**
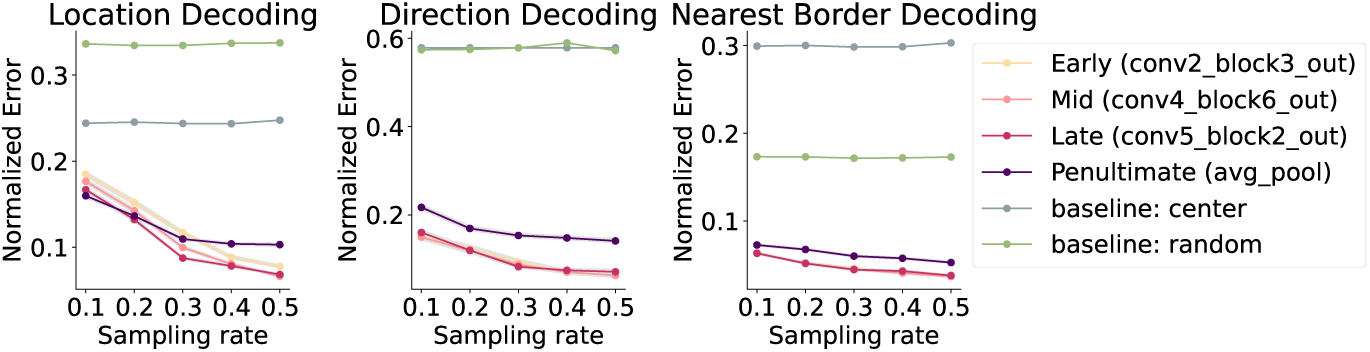
ResNet-50 (untrained). Similar to ResNet-50 pretrained on images, the untrained counterpart can effectively decode spatial knowledge related to location, heading direction and distance to borders. All layers decoder better than the baseline decoders which do not rely on visual signals of the environment.

**Figure S4:**
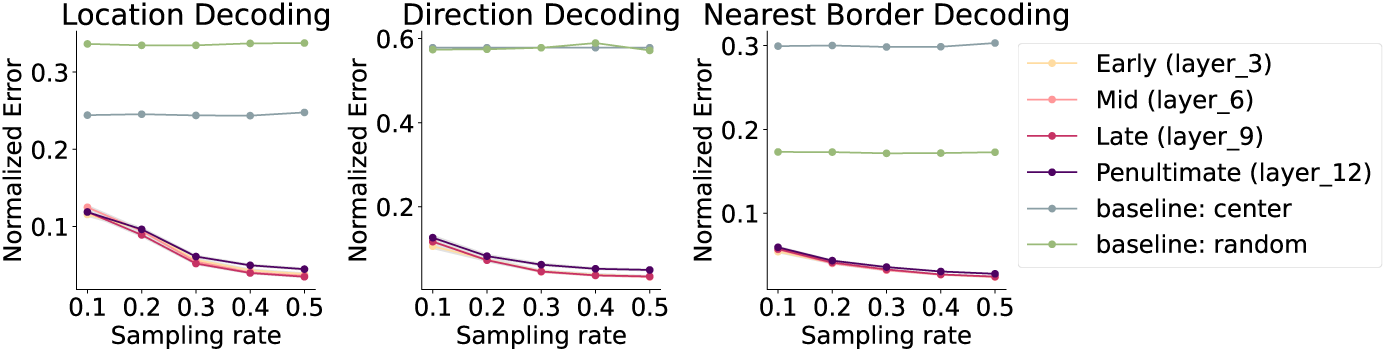
ViT-B/16 (trained). Linear decoders trained on different layers of the pretrained ViT model show very similar decoding performance. Overall the decoding performance is much better than the two baseline decoders which do not incorporate visual signals.

**Figure S5:**
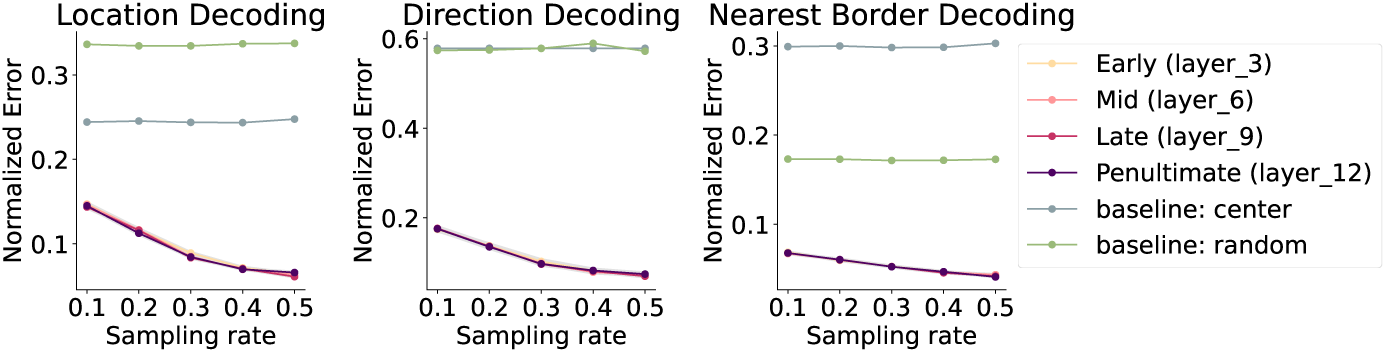
ViT-B/16 (untrained). Linear decoders trained on the untrained ViT model also achieve accurate decoding performance on location, heading direction and distance to the nearest border. Similar to the trained version, all layers considered in our analysis achieve comparable performance and outperform the baselines.

## **B** Distribution of cell types of spatial units across models and layers

**Figure S6:**
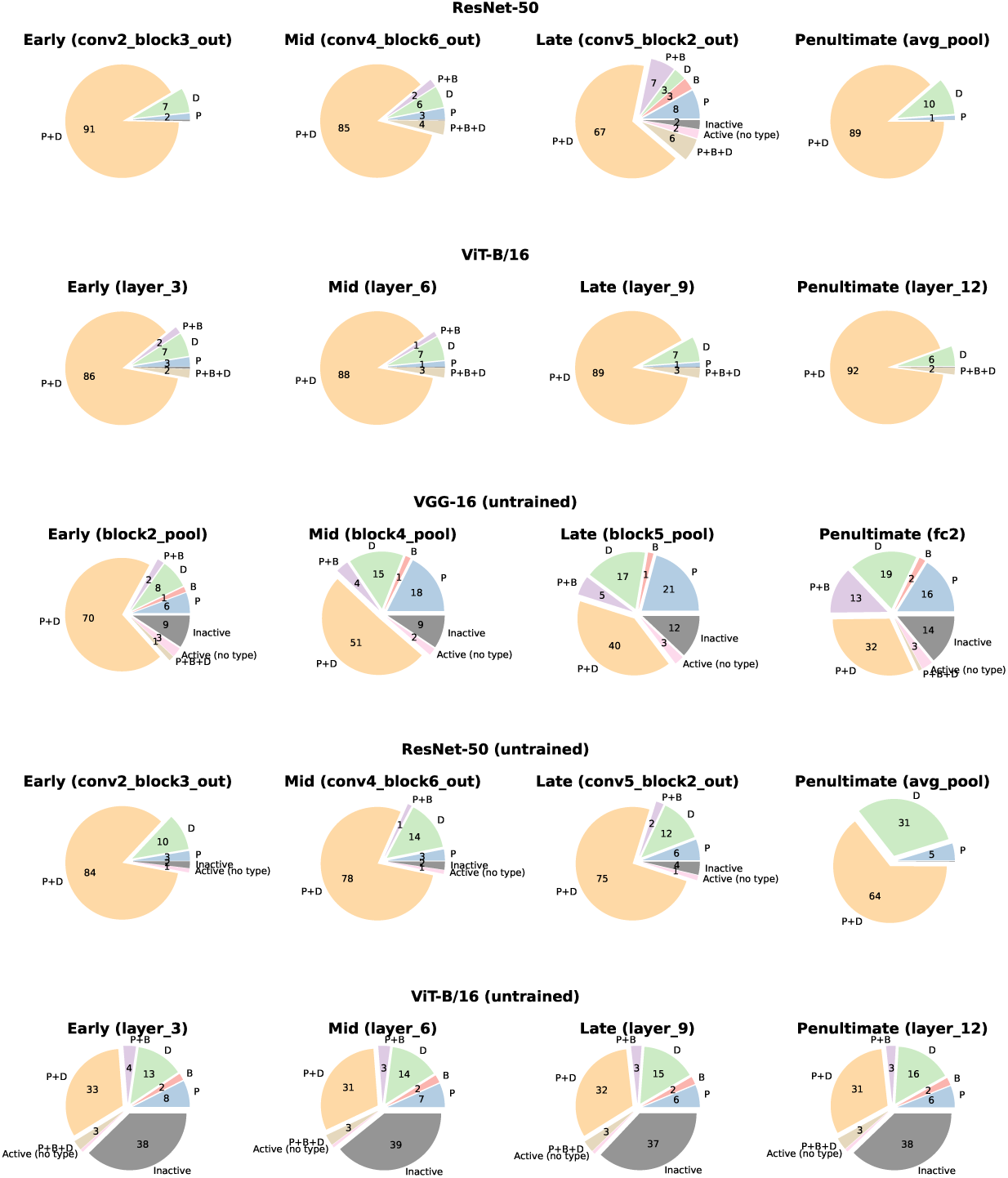
**Distribution of different spatial unit types across layers of perceptual models of object recognition.**

## **C** Model units with spatial firing profiles

**Figure S7:**
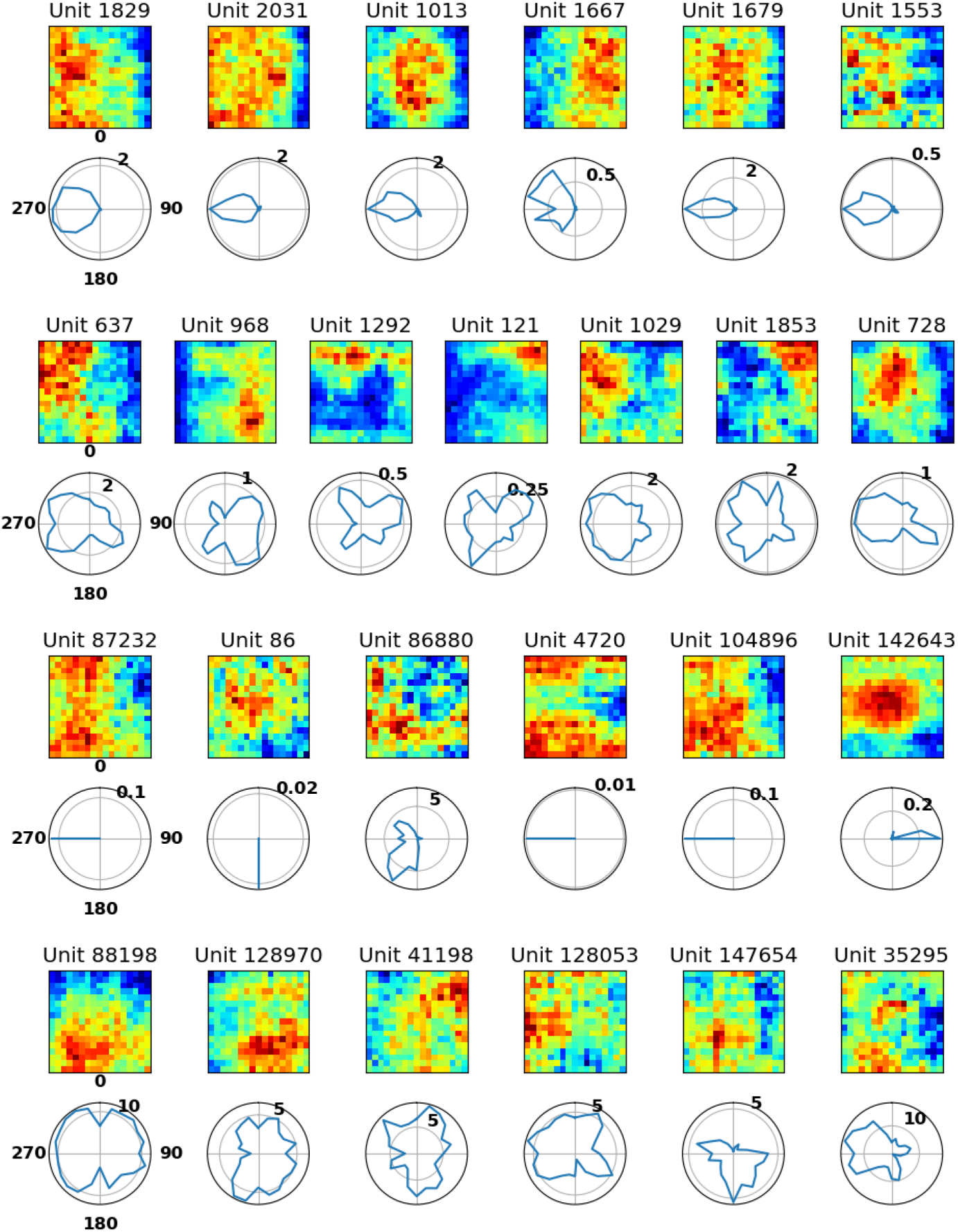
**Examples of model units exhibiting spatial characteristics.**

## **D** Spatial decoding performance under unit exclusion across various perception models

**Figure S8:**
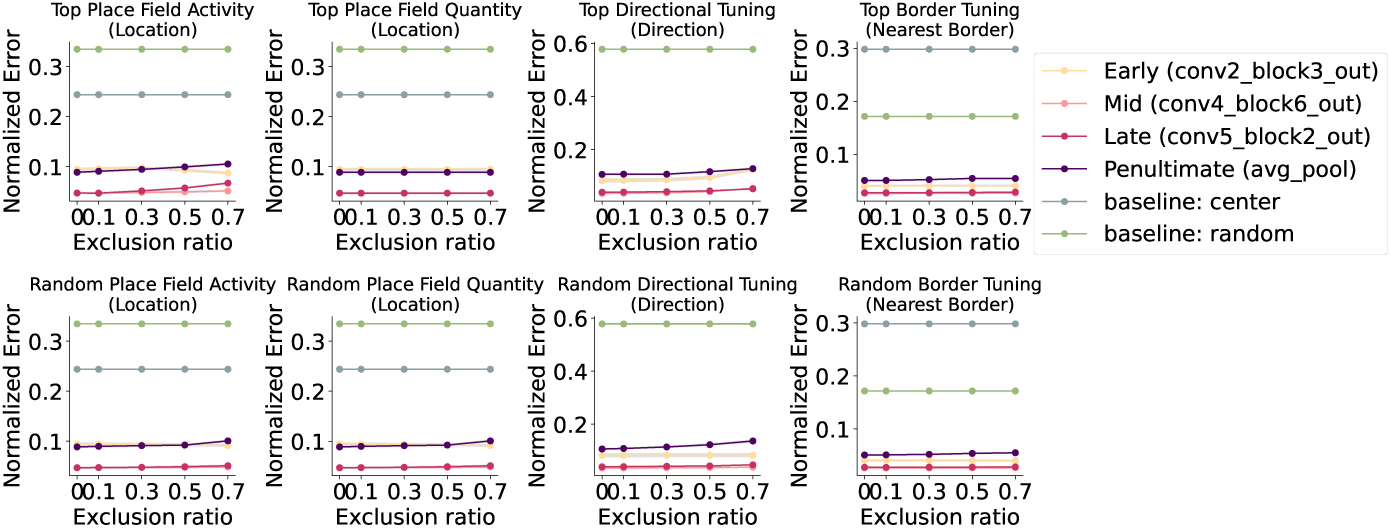
**Excluding units with the strongest spatial profiles had minimal impact on spatial knowledge (ResNet-50).**

**Figure S9:**
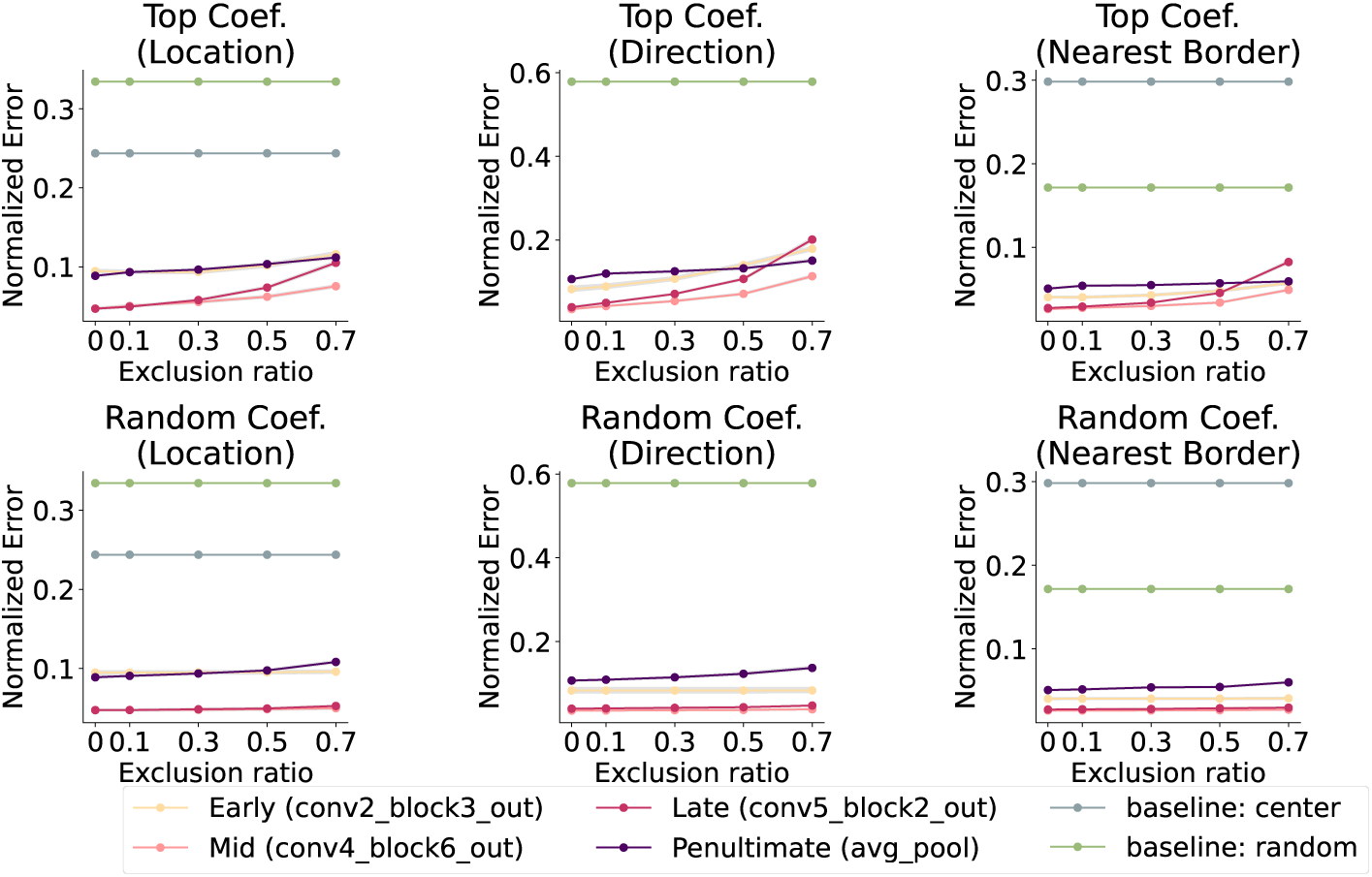
**Excluding units by task-relevance affects spatial decoding performance (ResNet-50).**

**Figure S10:**
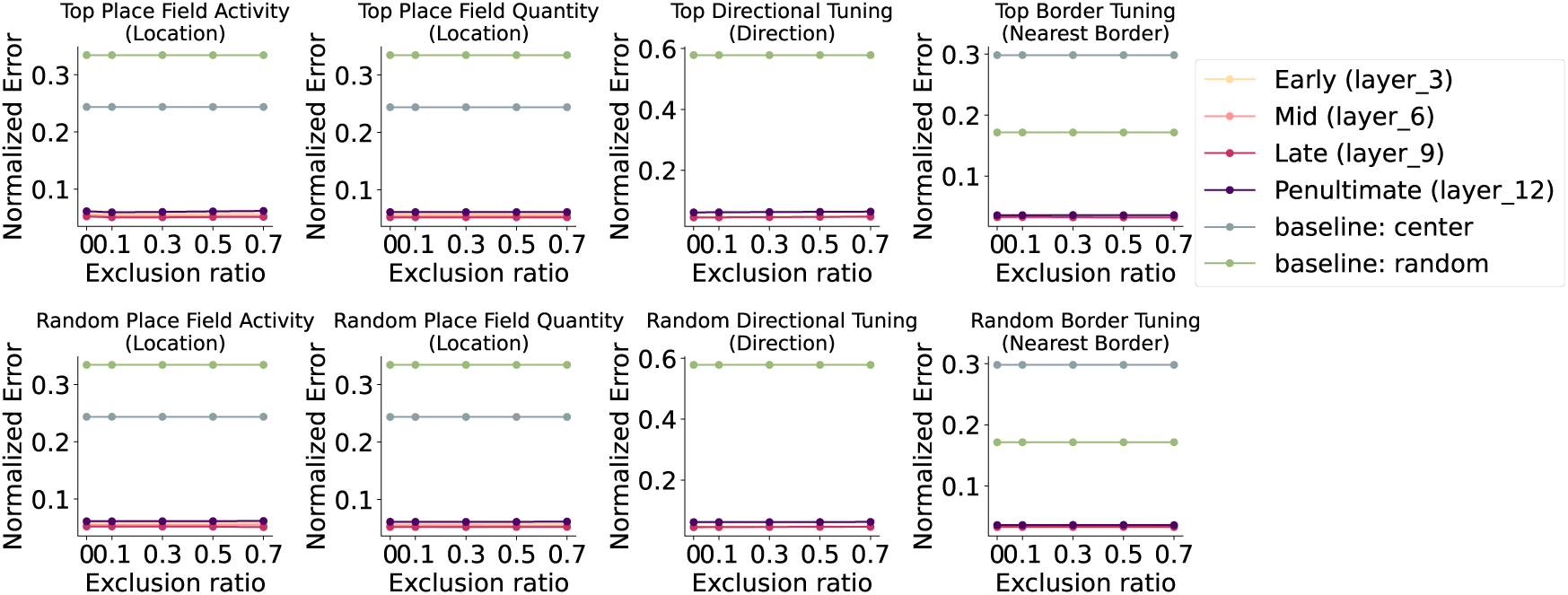
**Excluding units with the strongest spatial profiles had minimal impact on spatial knowledge (ViT-B/16).**

**Figure S11:**
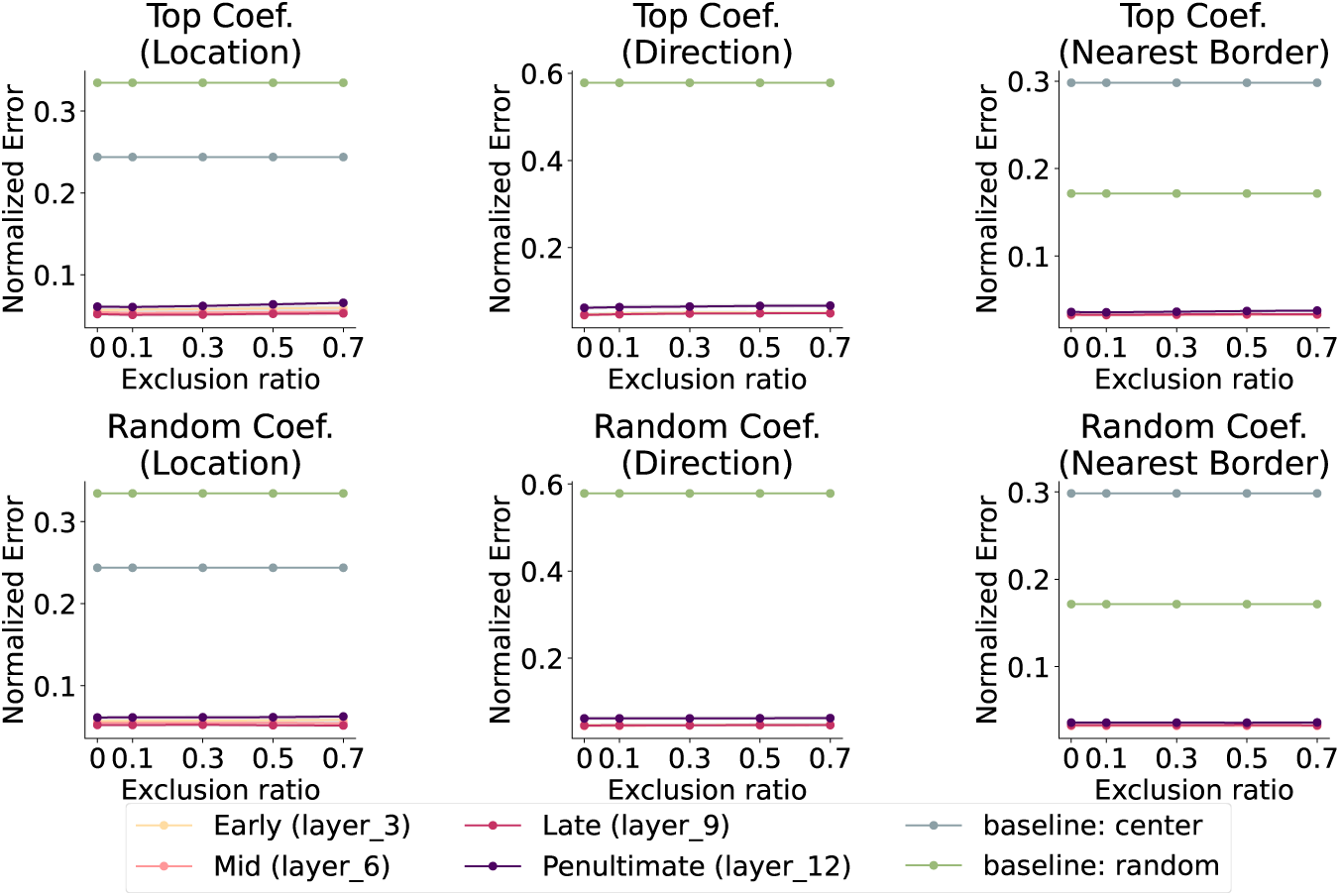
**Excluding units by task-relevance affects spatial decoding performance (ViT-B/16).**

## **E** Relationship between model units’ spatial properties and corresponding decoder coefficient strengths

**Figure S12:**
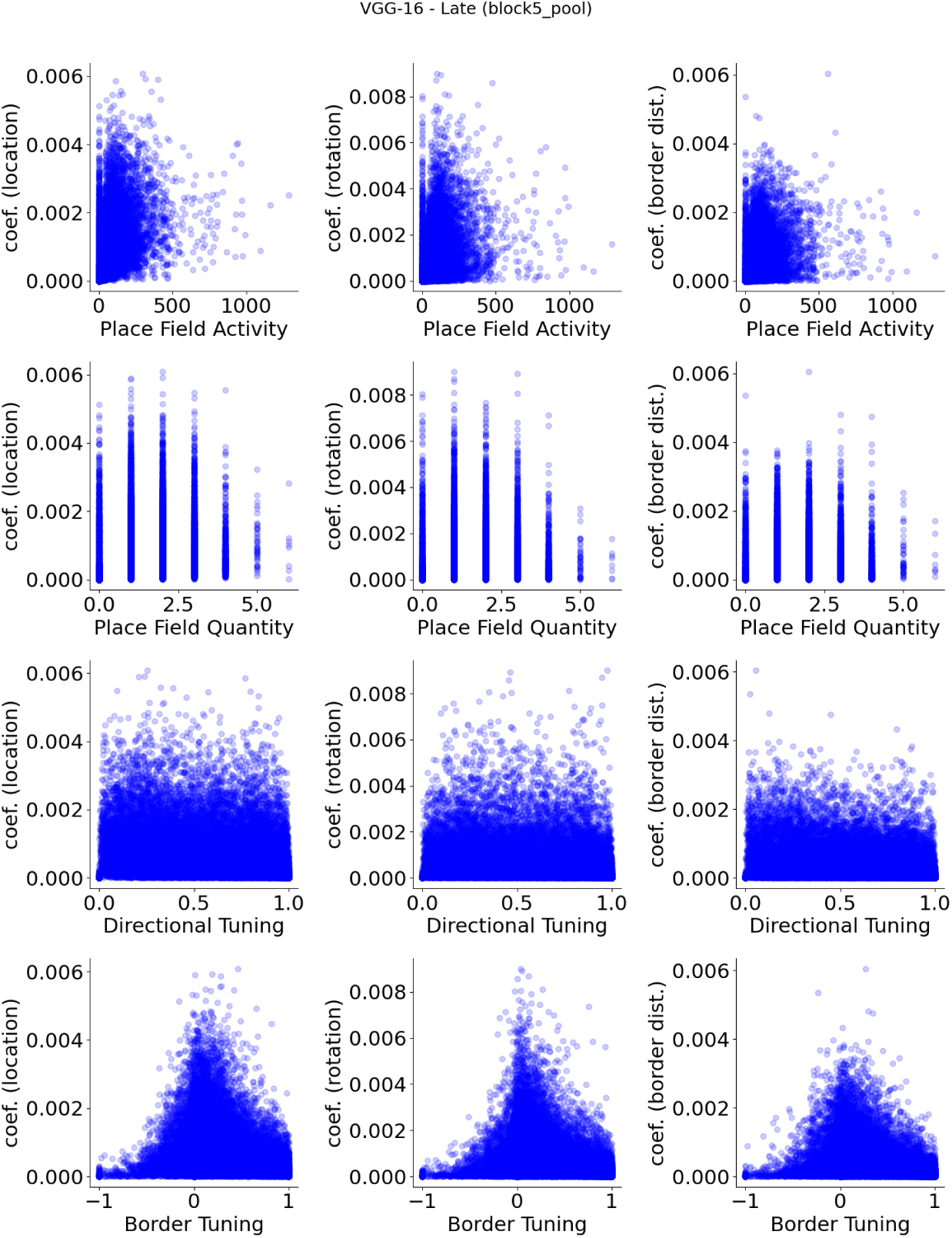
**VGG-16 (pre-trained), late layer (block5 pool) units, show no apparent relationship between spatial properties and their decoder weight strengths.**

**Figure S13:**
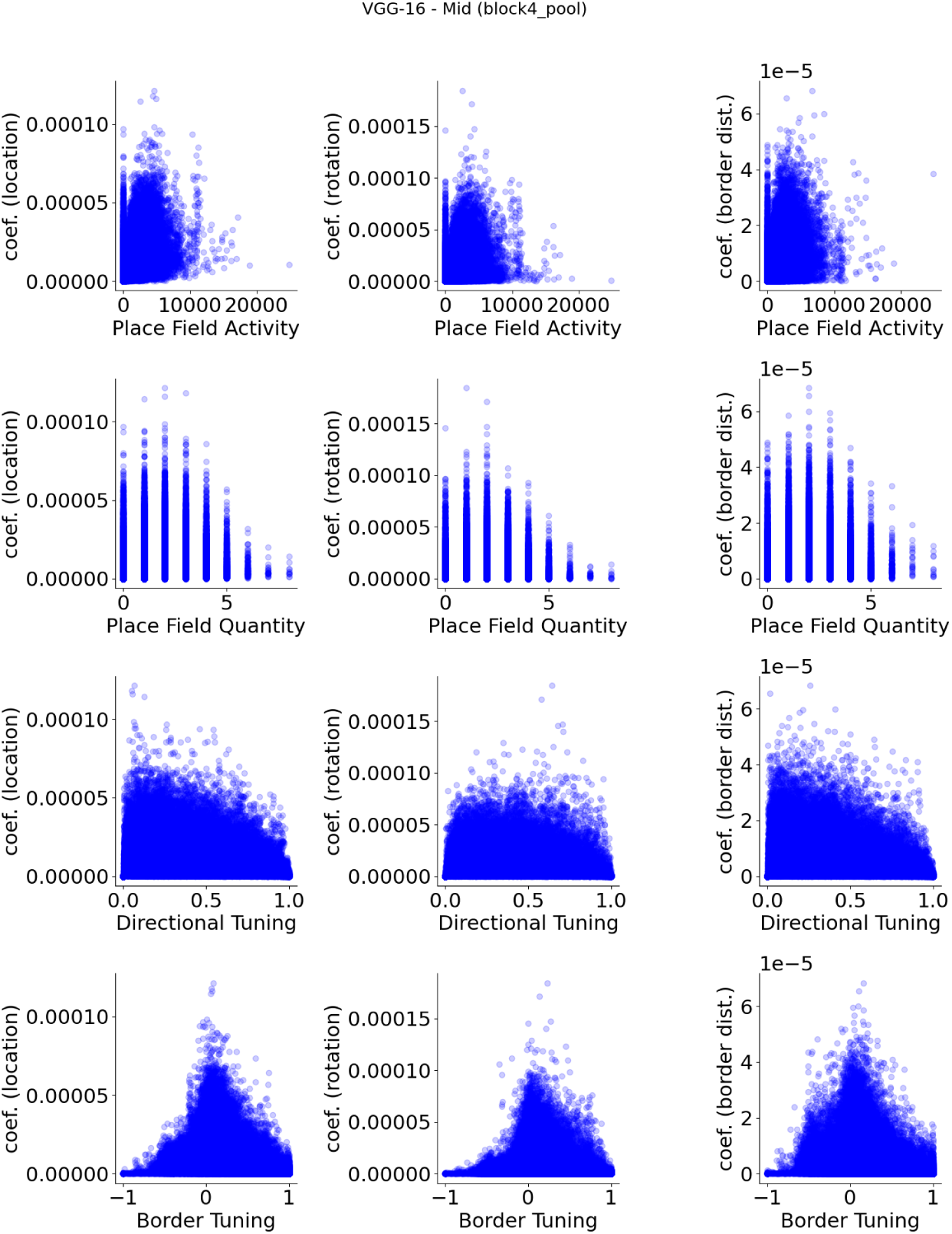
**VGG-16 (pre-trained), mid layer (block4 pool) units, show no apparent relationship between spatial properties and their decoder weight strengths.**

**Figure S14:**
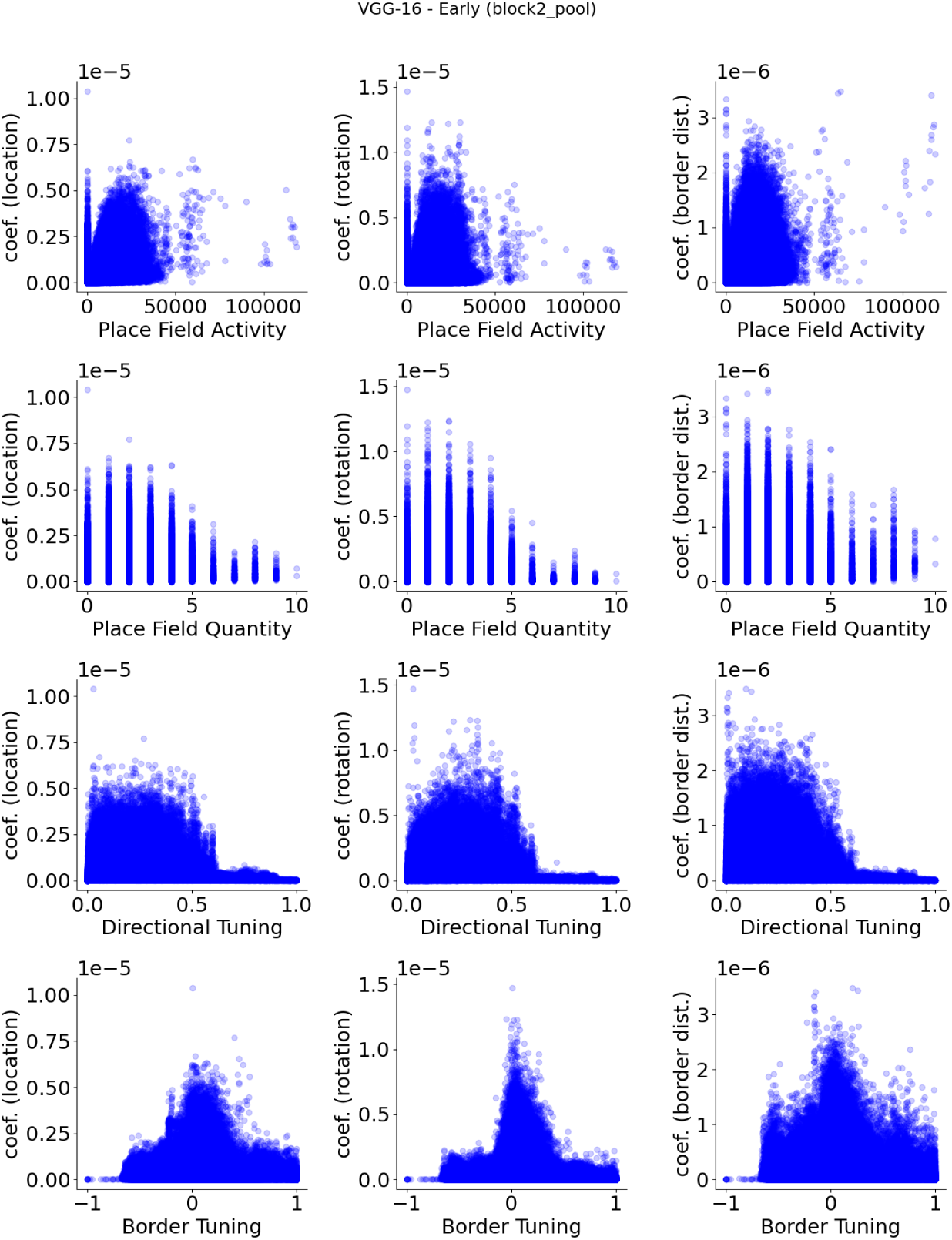
**VGG-16 (pre-trained), early layer (block2 pool) units, show no apparent relationship between spatial properties and their decoder weight strengths.**

**Figure S15:**
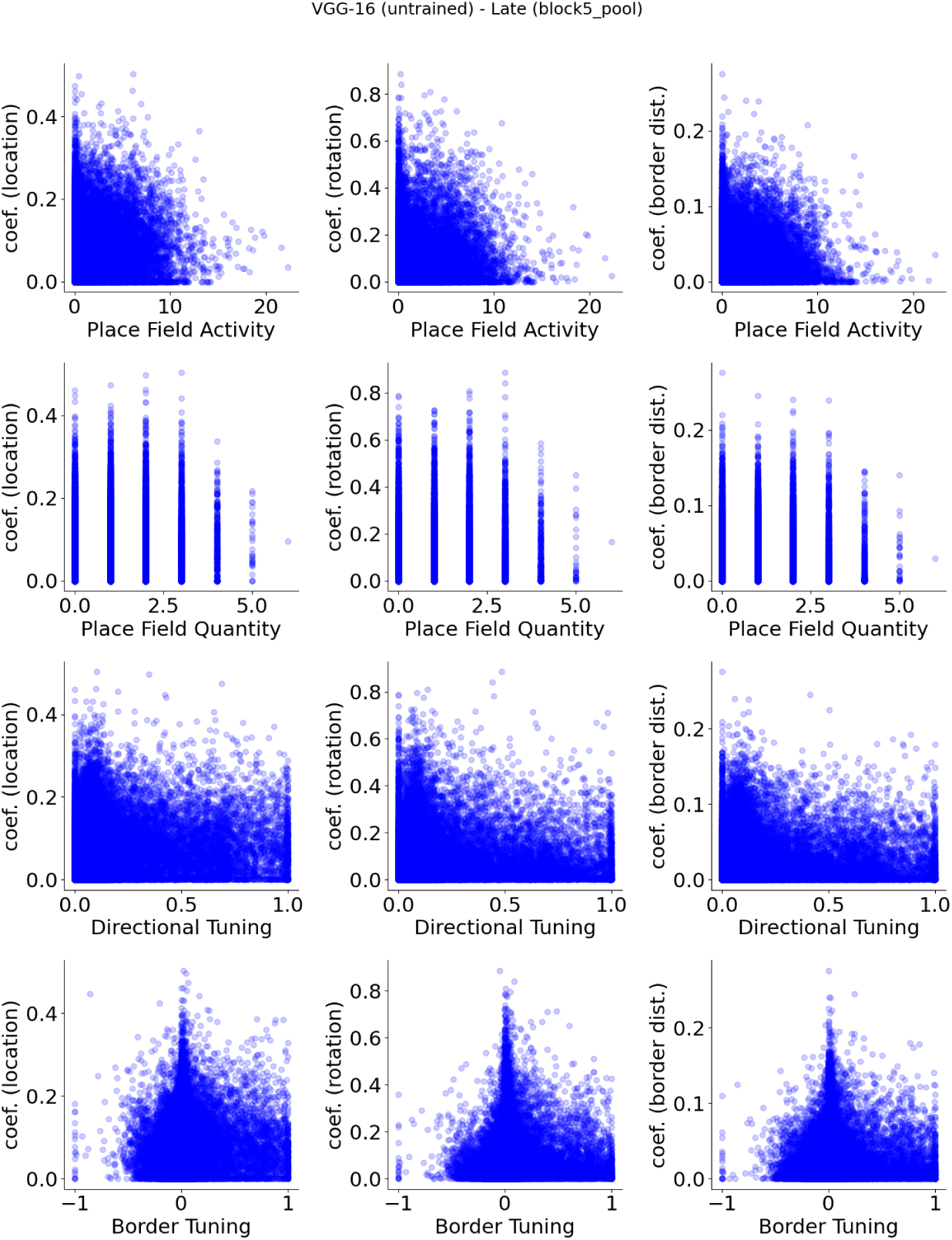
**VGG-16 (untrained), late layer (block5 pool) units, show no apparent relationship between spatial properties and their decoder weight strengths.**

**Figure S16:**
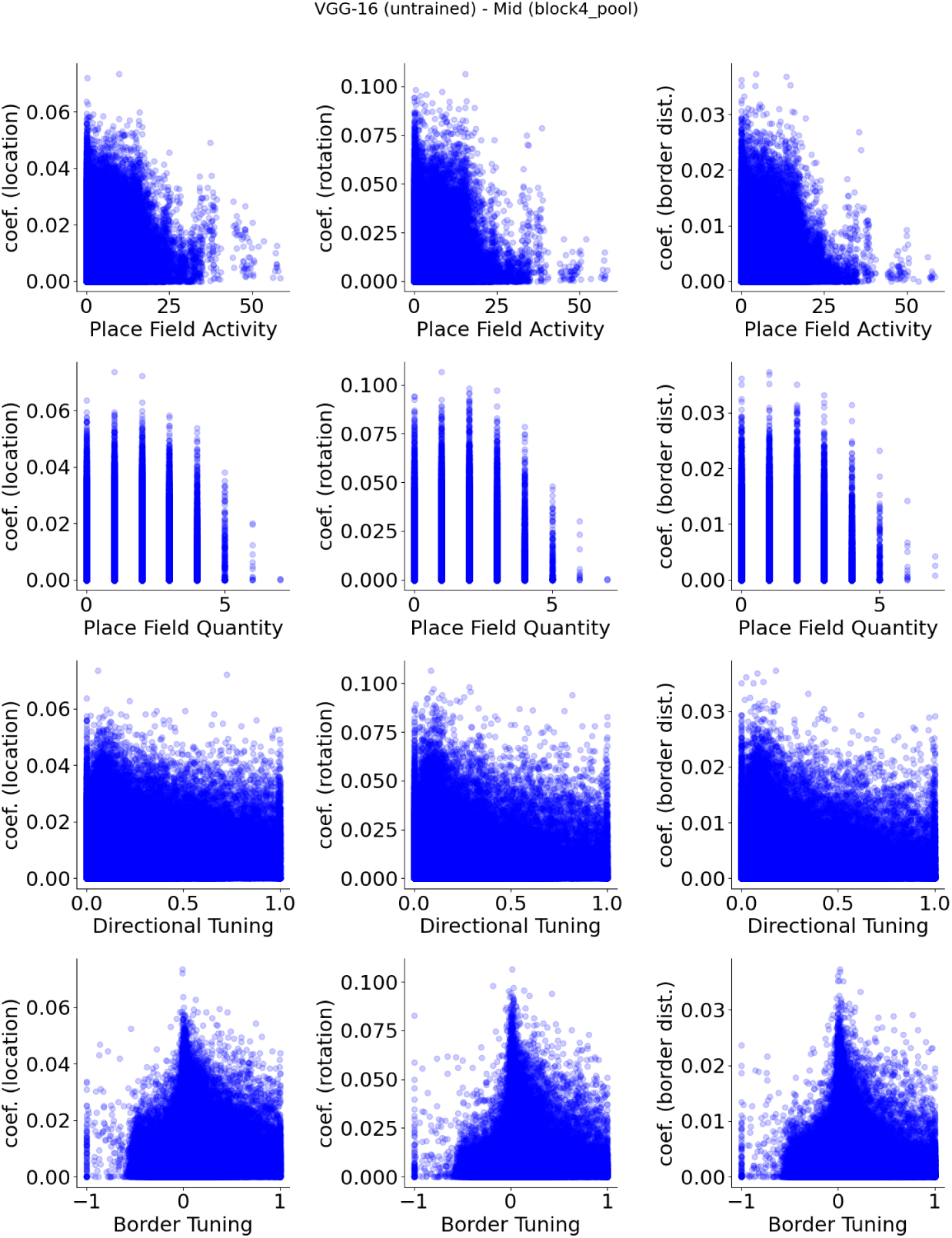
**VGG-16 (untrained), mid layer (block4 pool) units, show no apparent relationship between spatial properties and their decoder weight strengths.**

**Figure S17:**
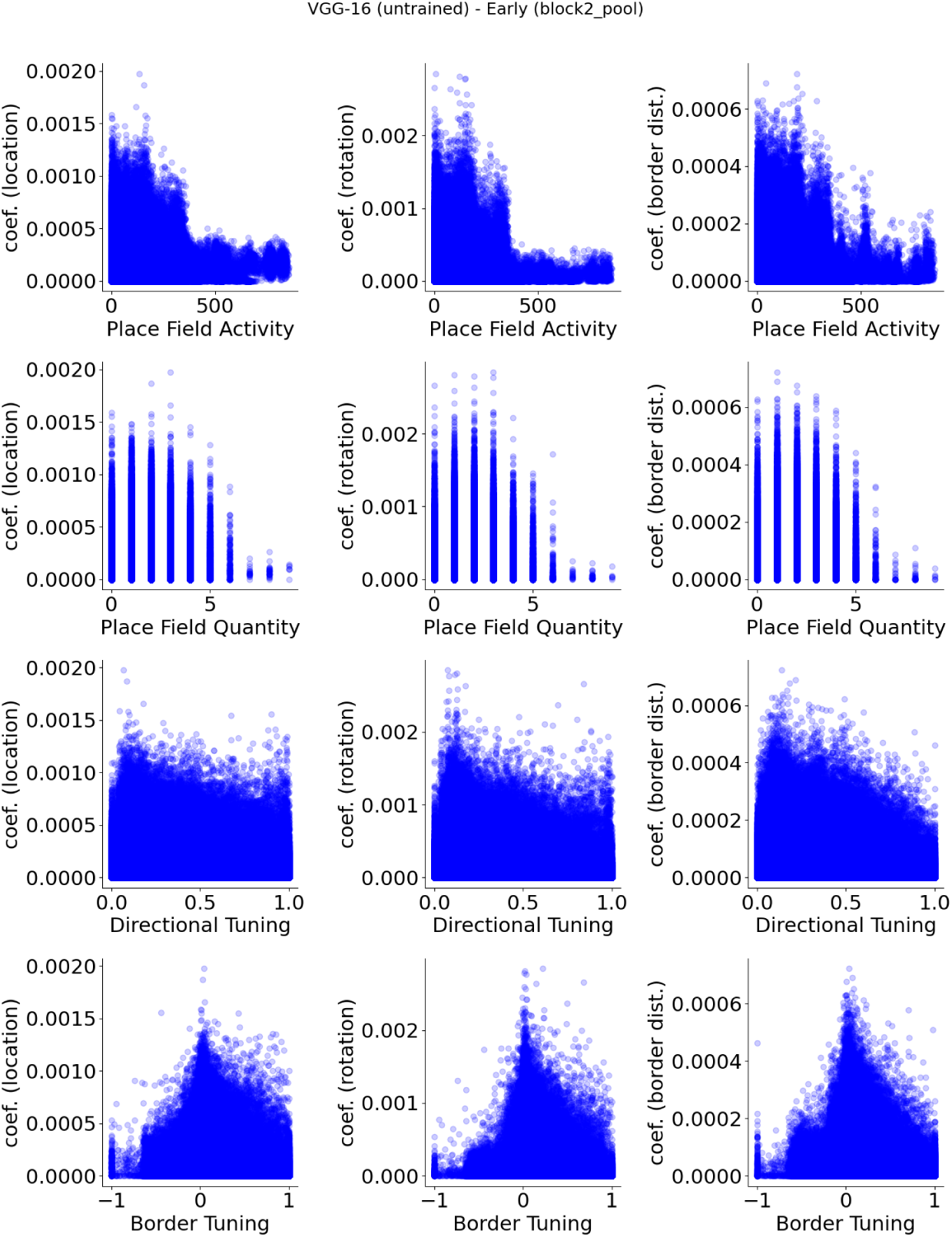
**VGG-16 (untrained), early layer (block2 pool) units, show no apparent relationship between spatial properties and their decoder weight strengths.**

**Figure S18:**
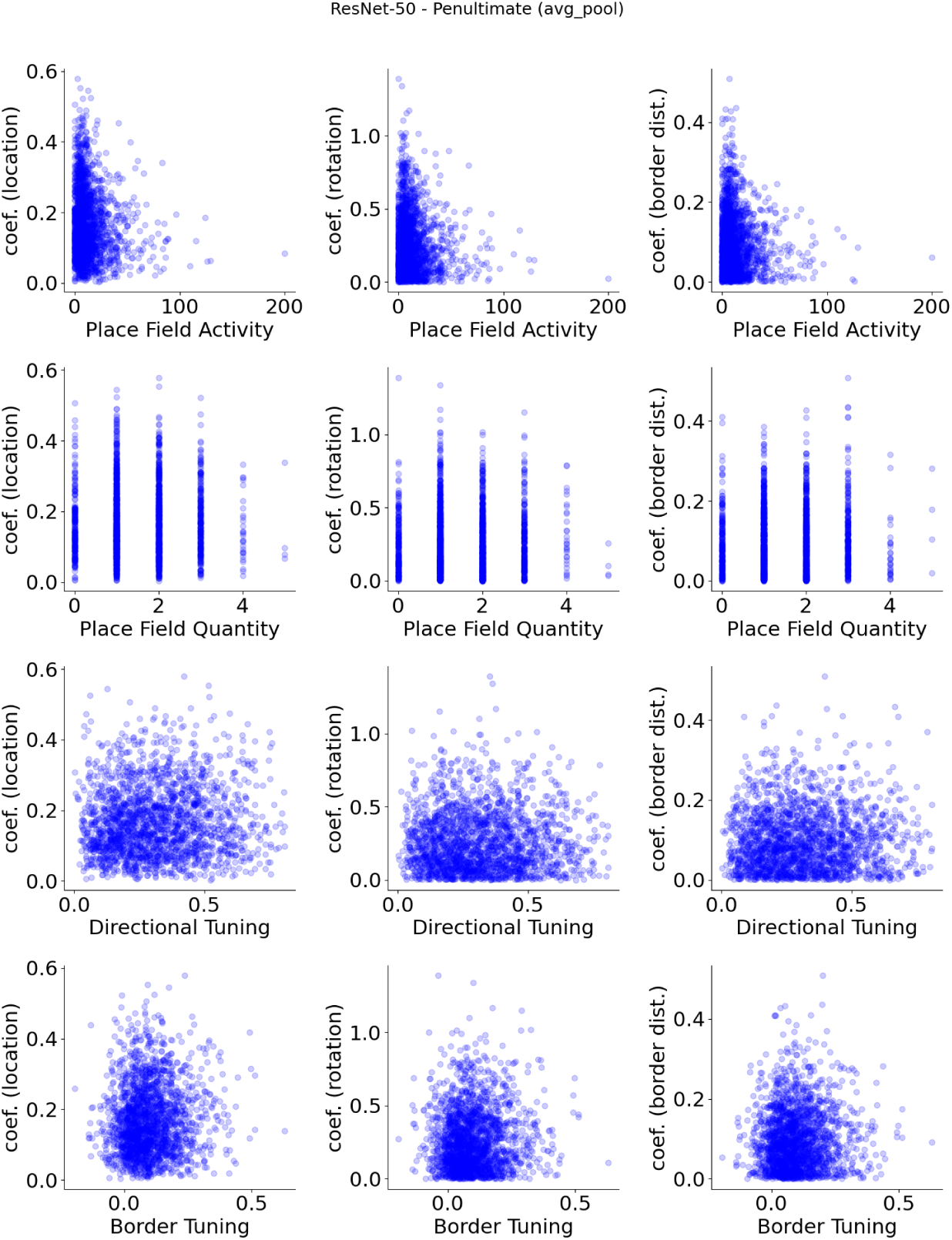
**ResNet-50 (pre-trained), penultimate layer (avg pool) units, show no apparent relationship between spatial properties and their decoder weight strengths.**

**Figure S19:**
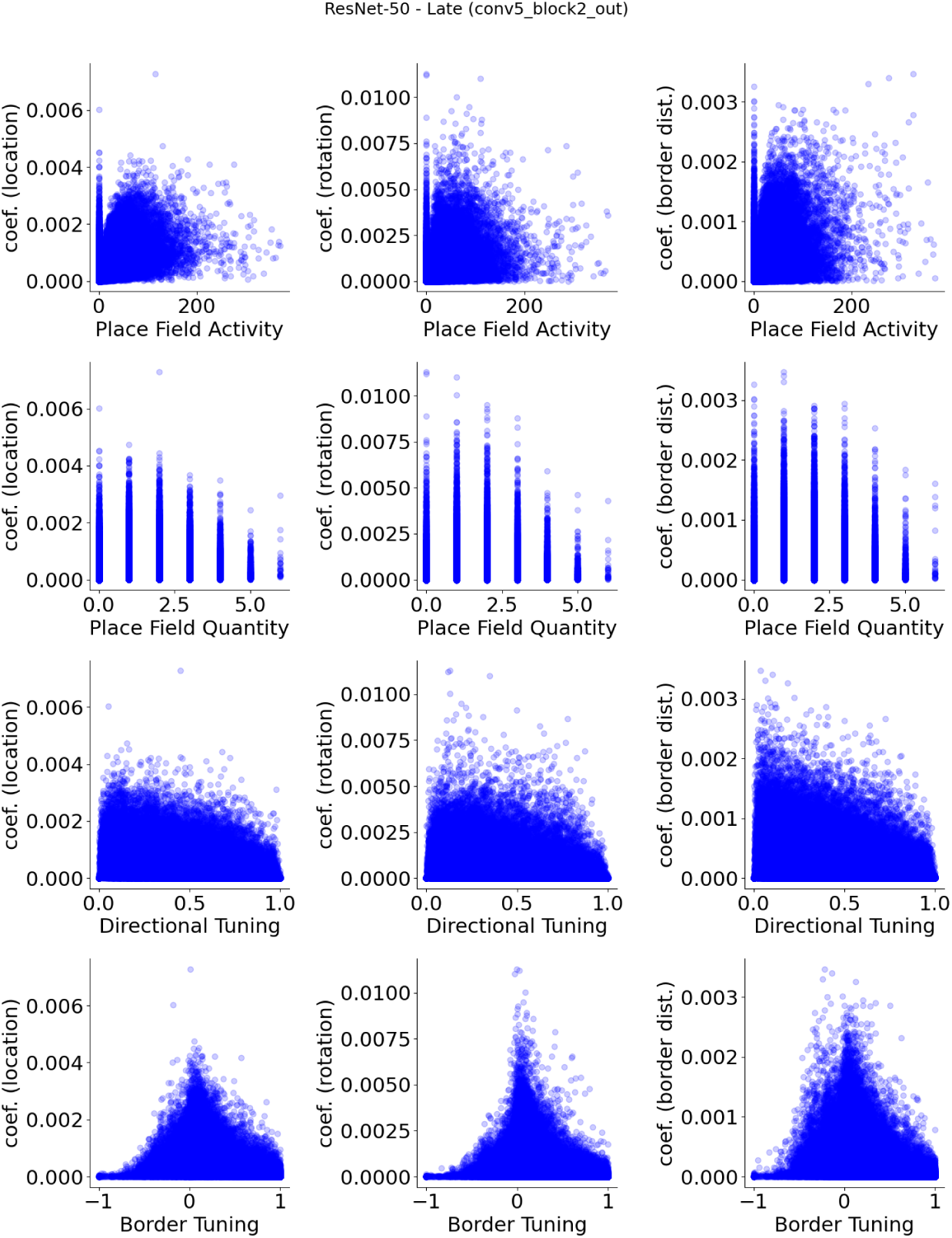
**ResNet-50 (pre-trained), late layer (conv5 block2) units, show no apparent relationship between spatial properties and their decoder weight strengths.**

**Figure S20:**
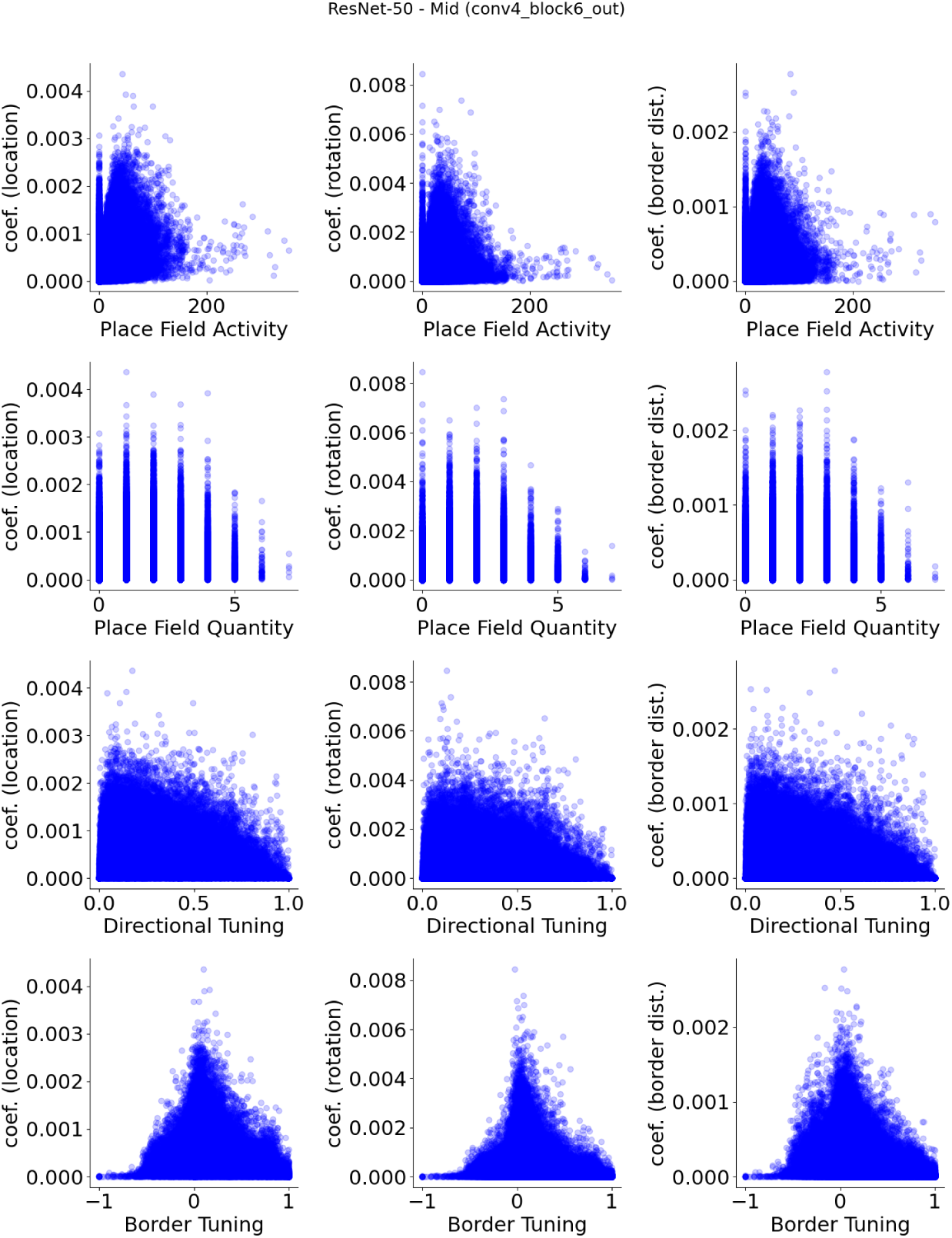
**ResNet-50 (pre-trained), mid layer (conv4 block6) units, show no apparent relationship between spatial properties and their decoder weight strengths.**

**Figure S21:**
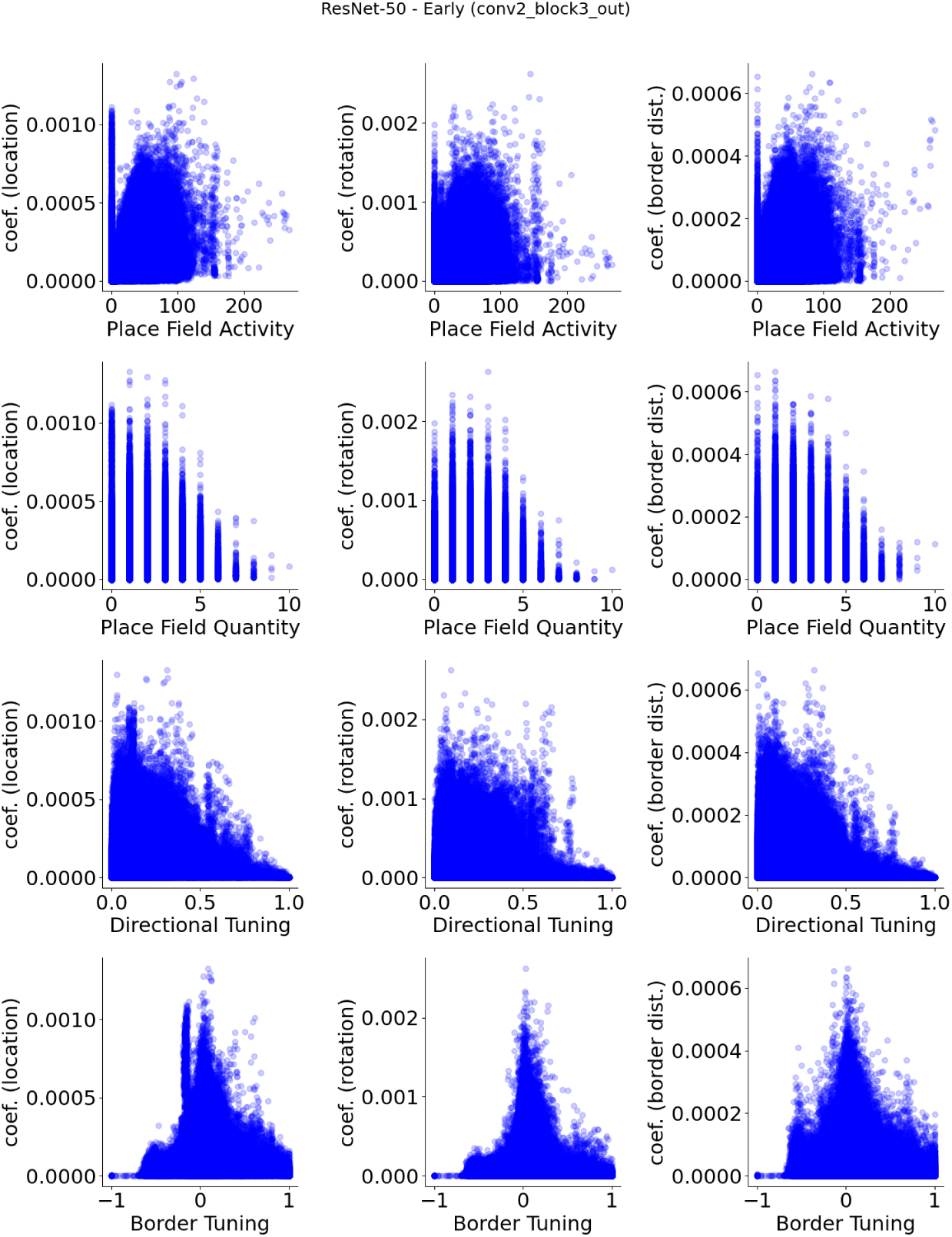
**ResNet-50 (pre-trained), early layer (conv2 block3) units, show no apparent relationship between spatial properties and their decoder weight strengths.**

**Figure S22:**
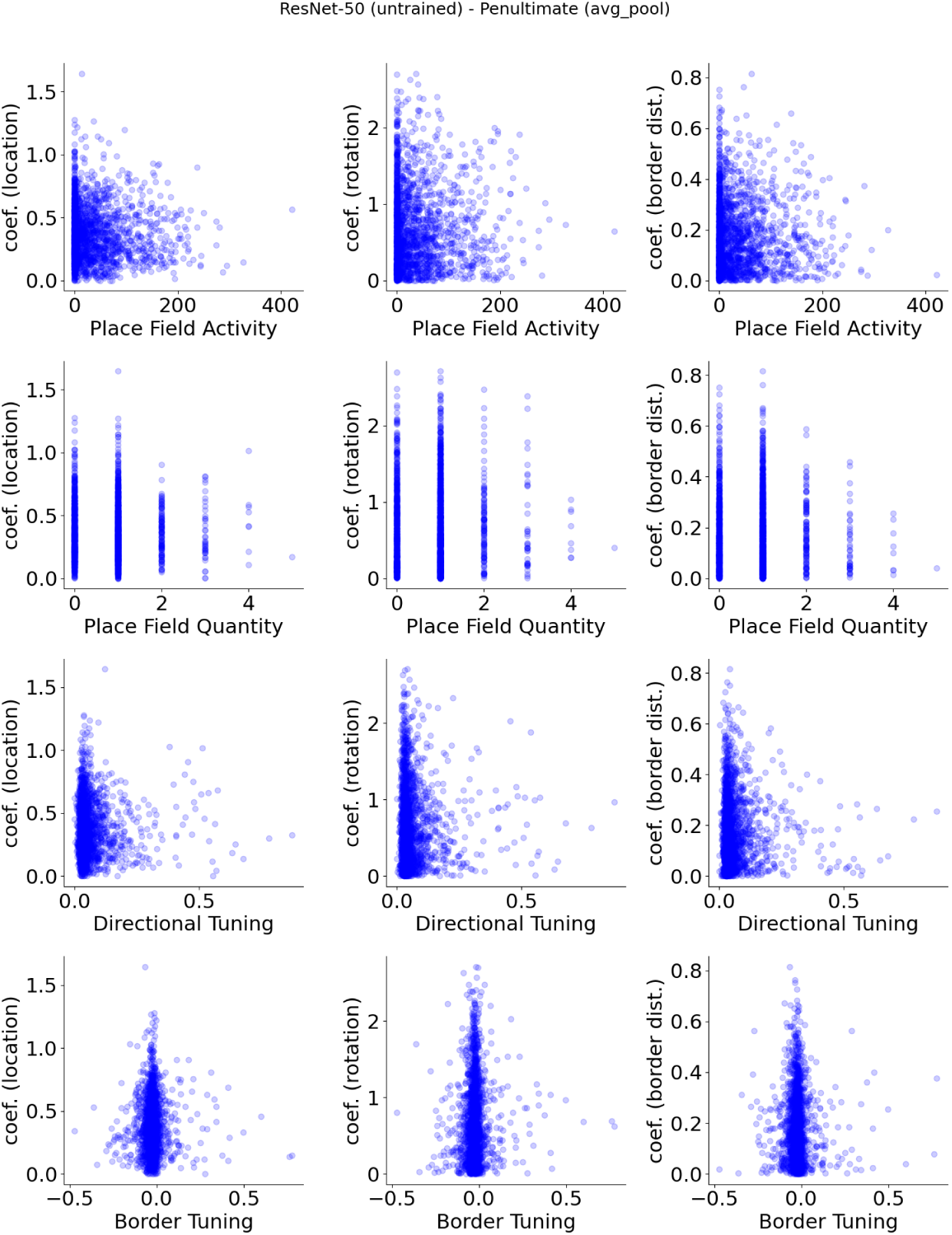
**ResNet-50 (untrained), penultimate layer (avg pool) units, show no apparent relationship between spatial properties and their decoder weight strengths.**

**Figure S23:**
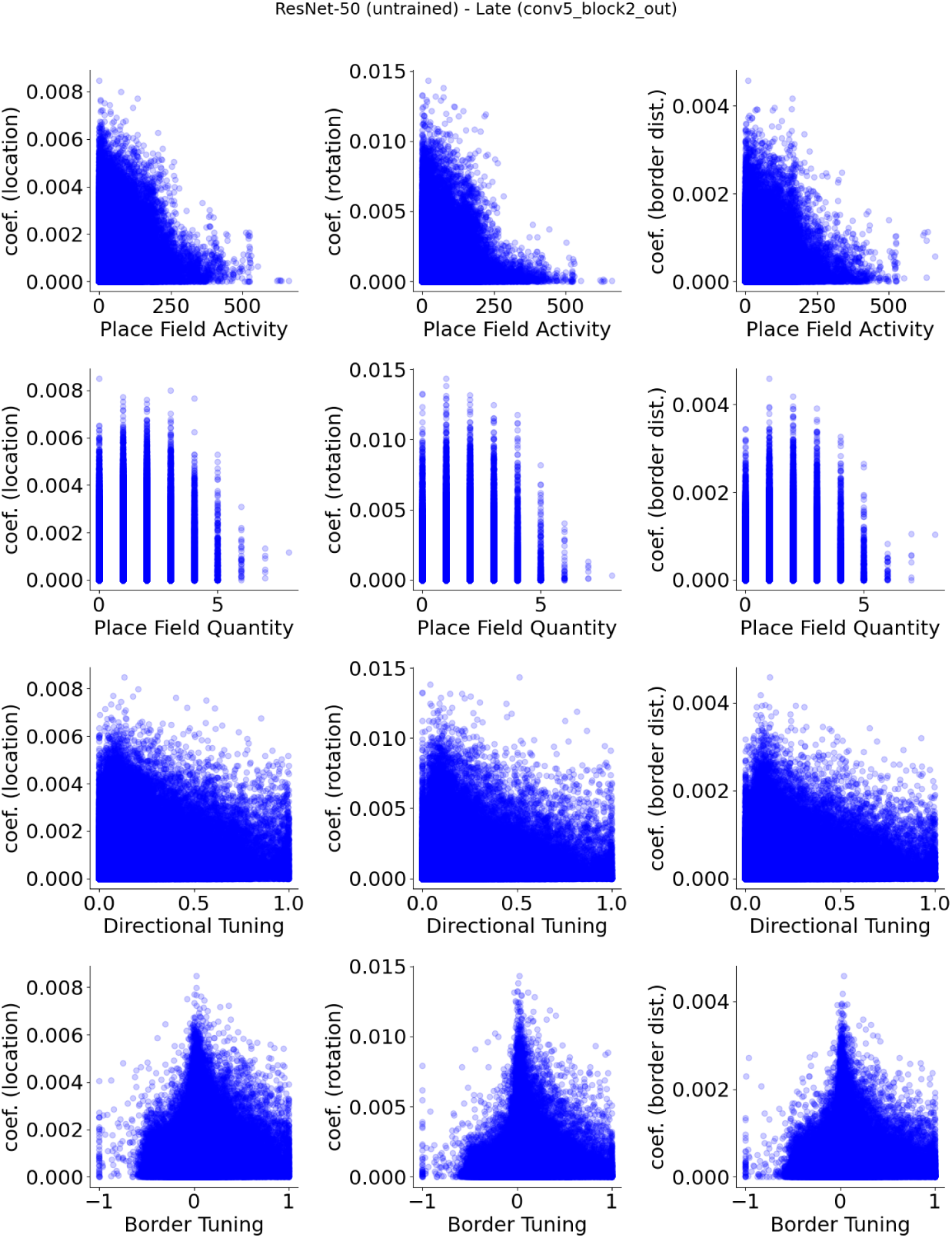
**ResNet-50 (untrained), late layer (conv5 block2) units, show no apparent relationship between spatial properties and their decoder weight strengths.**

**Figure S24:**
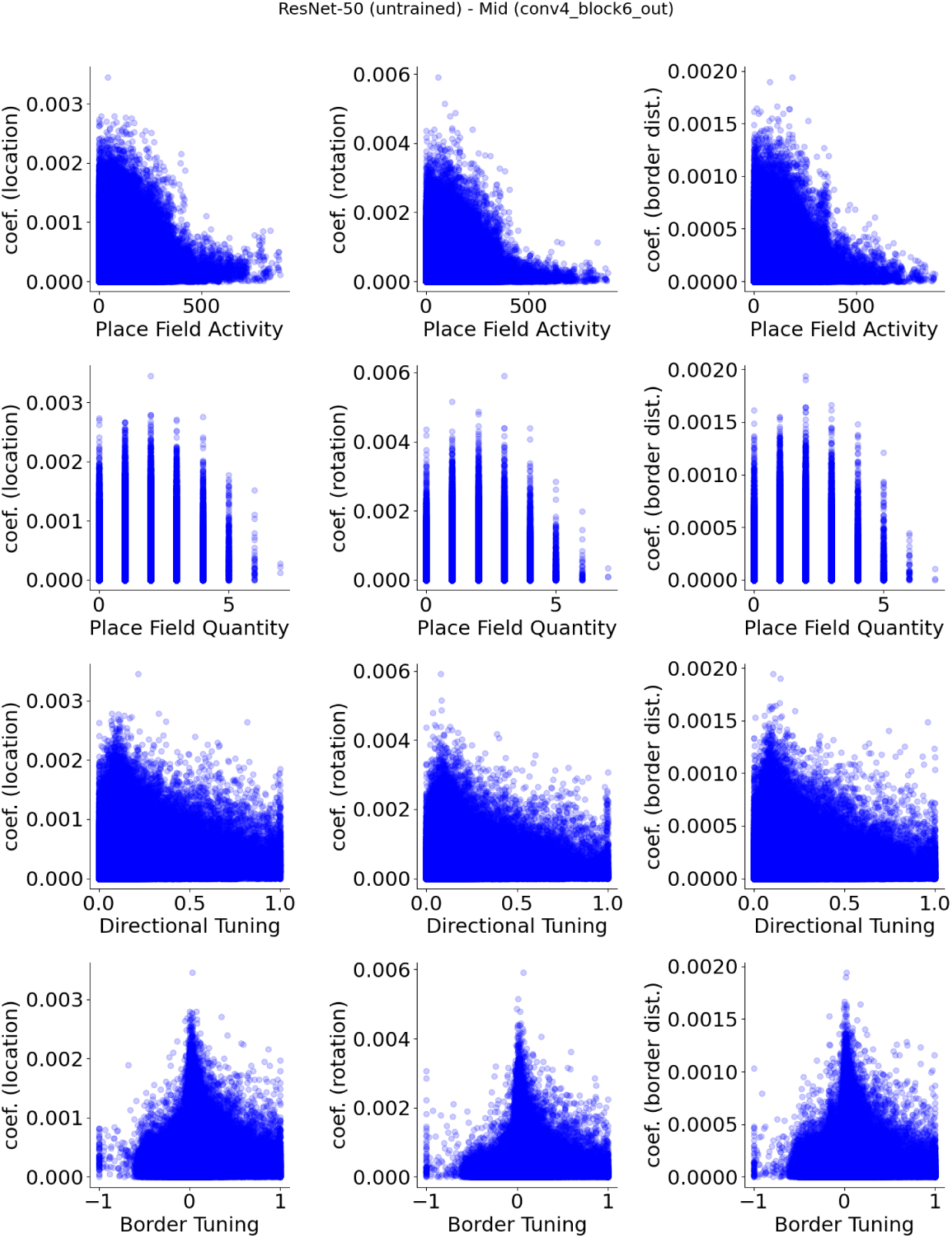
**ResNet-50 (untrained), mid layer (conv4 block6) units, show no apparent relationship between spatial properties and their decoder weight strengths.**

**Figure S25:**
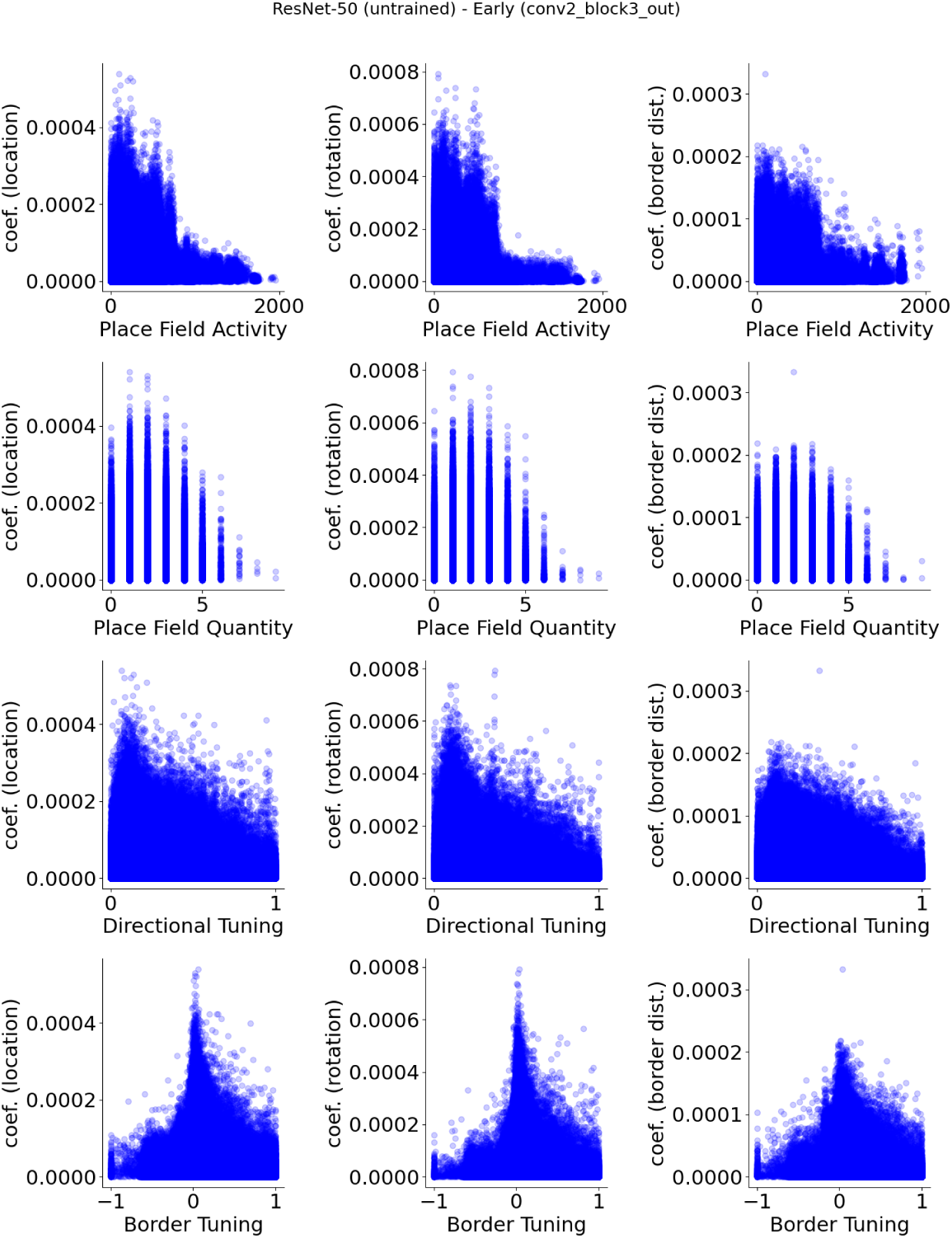
**ResNet-50 (untrained), early layer (conv2 block3) units, show no apparent relationship between spatial properties and their decoder weight strengths.**

**Figure S26:**
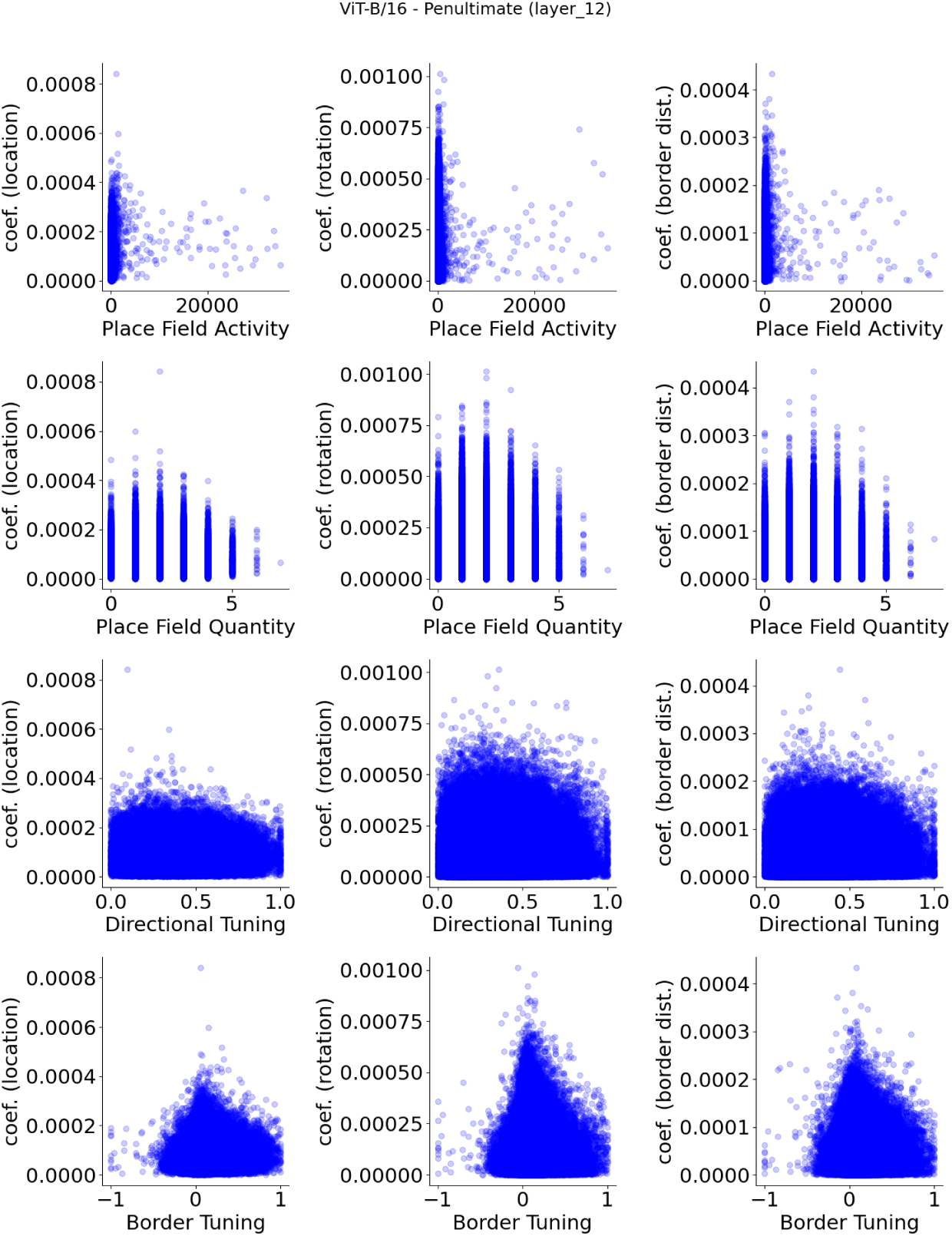
**ViT-B/16 (pre-trained), penultimate layer (layer 12) units, show no apparent relationship between spatial properties and their decoder weight strengths.**

**Figure S27:**
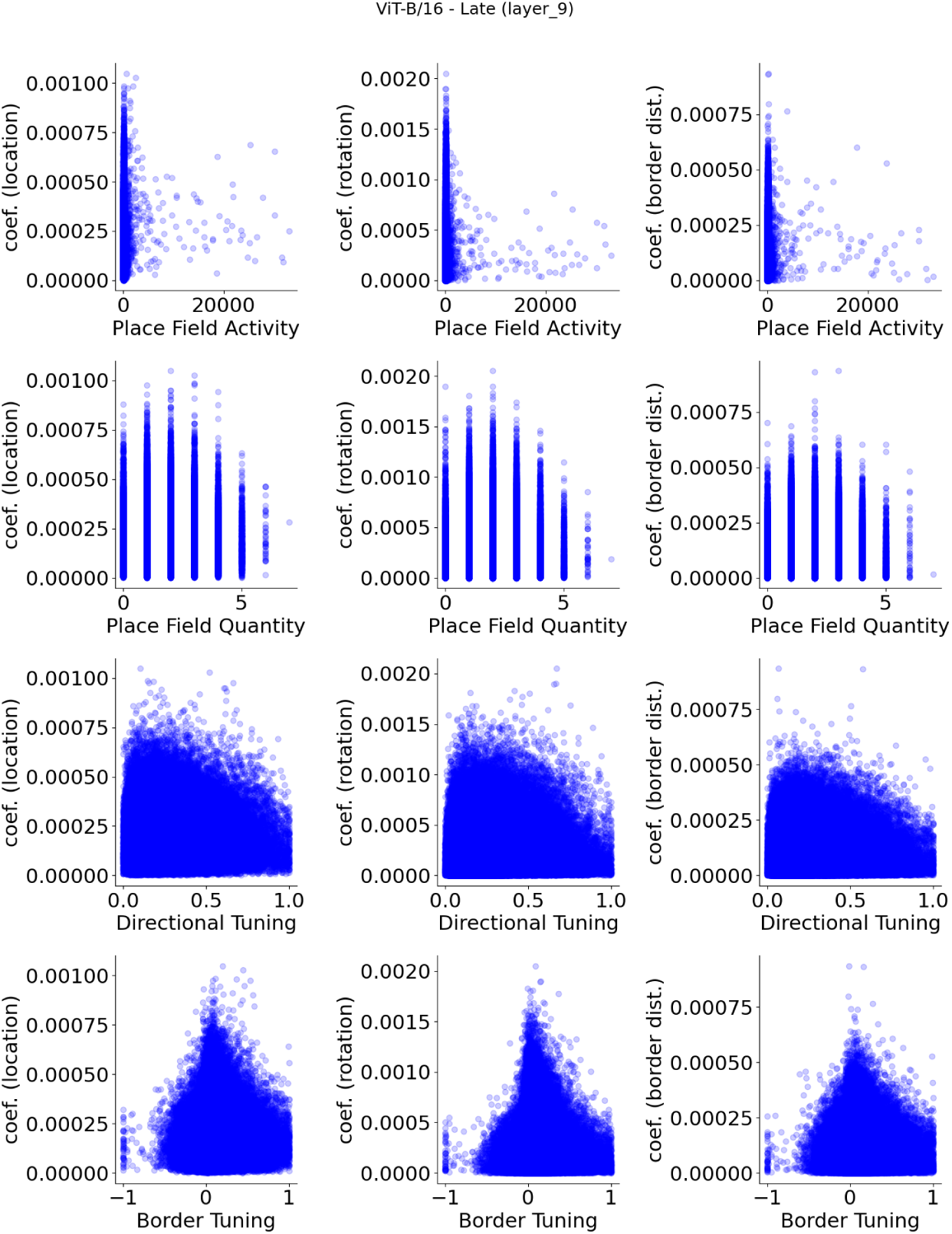
**ViT-B/16 (pre-trained), late layer (layer 9) units, show no apparent relationship between spatial properties and their decoder weight strengths.**

**Figure S28:**
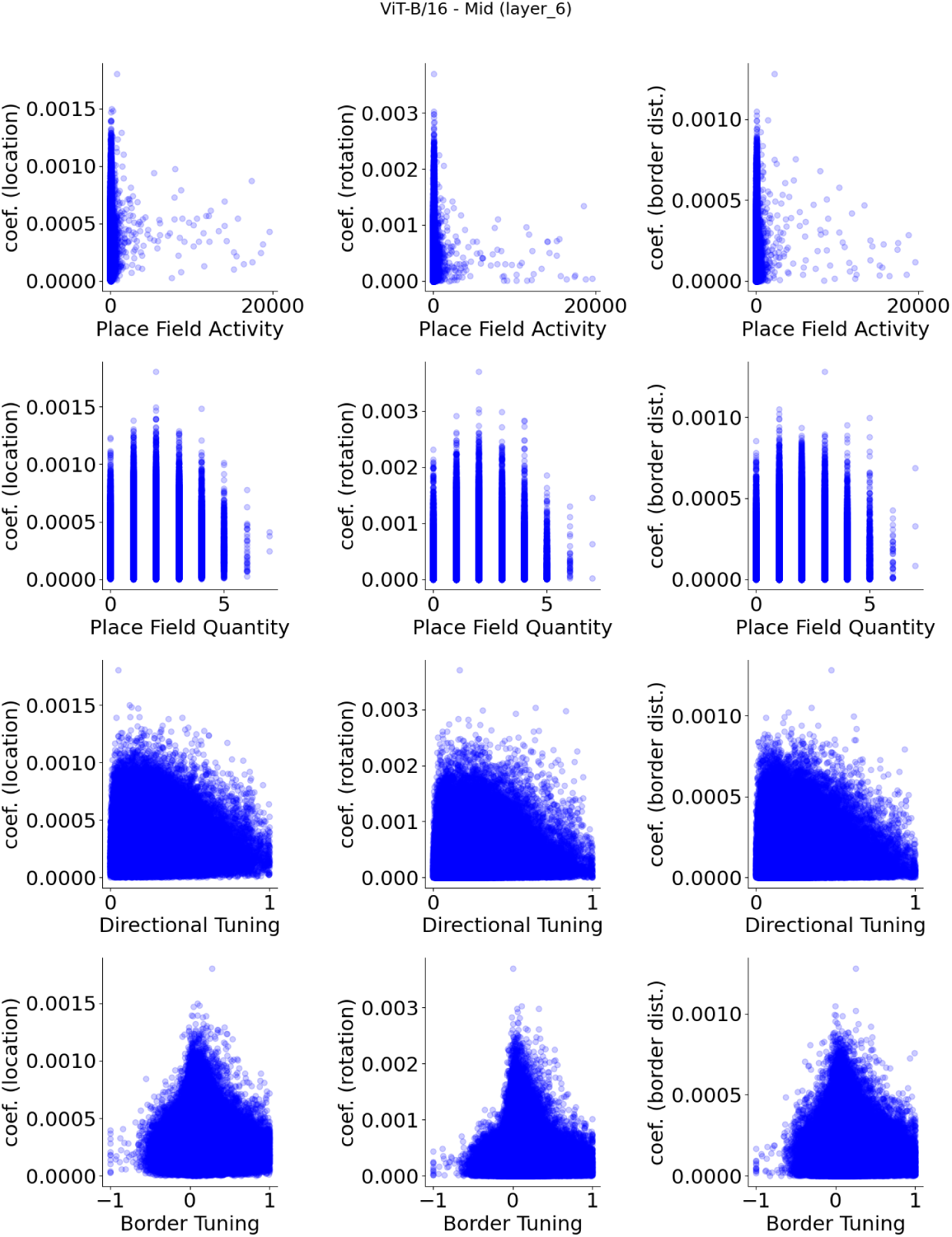
**ViT-B/16 (pre-trained), mid layer (layer 6) units, show no apparent relationship between spatial properties and their decoder weight strengths.**

**Figure S29:**
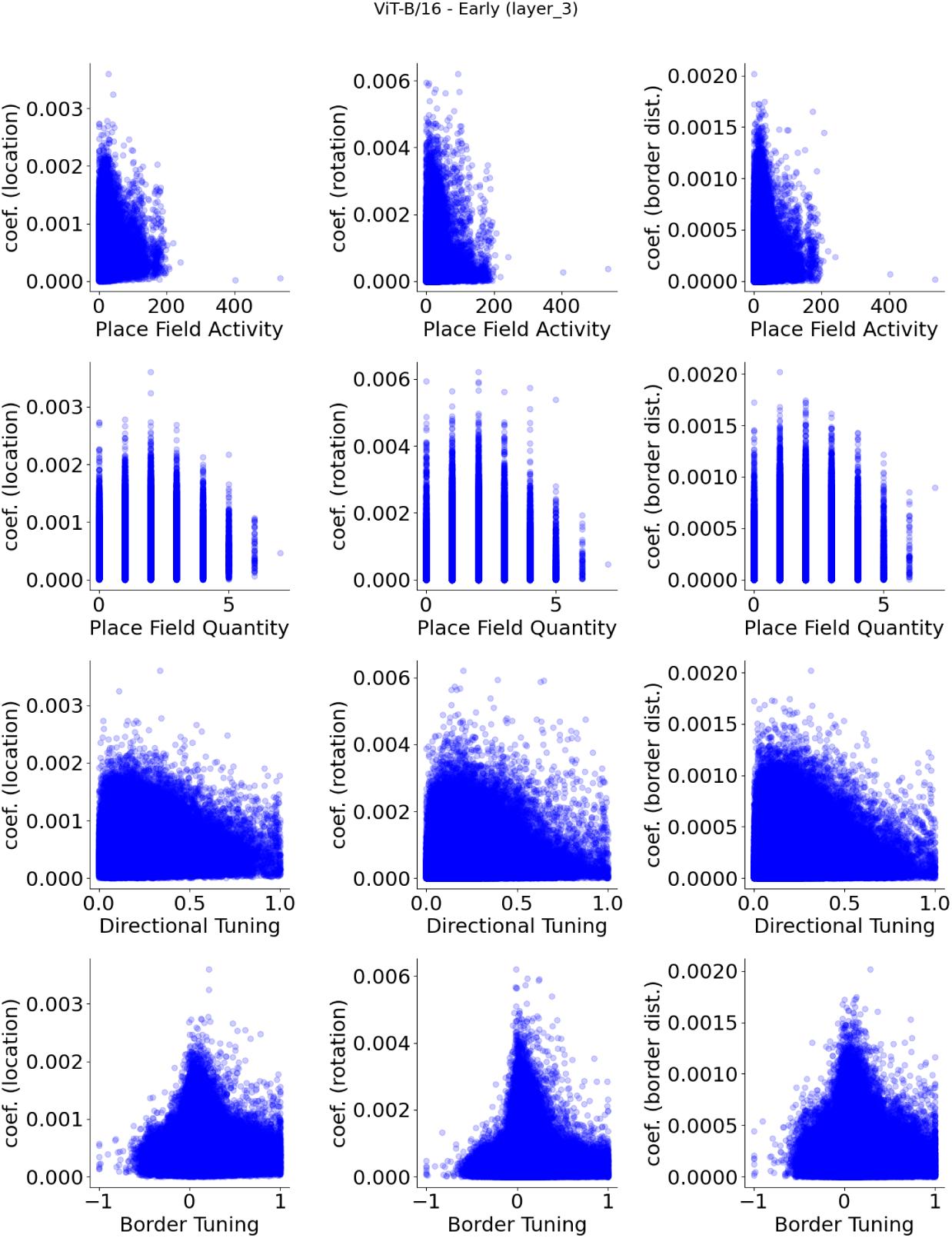
**ViT-B/16 (pre-trained), early layer (layer 3) units, show no apparent relationship between spatial properties and their decoder weight strengths.**

**Figure S30:**
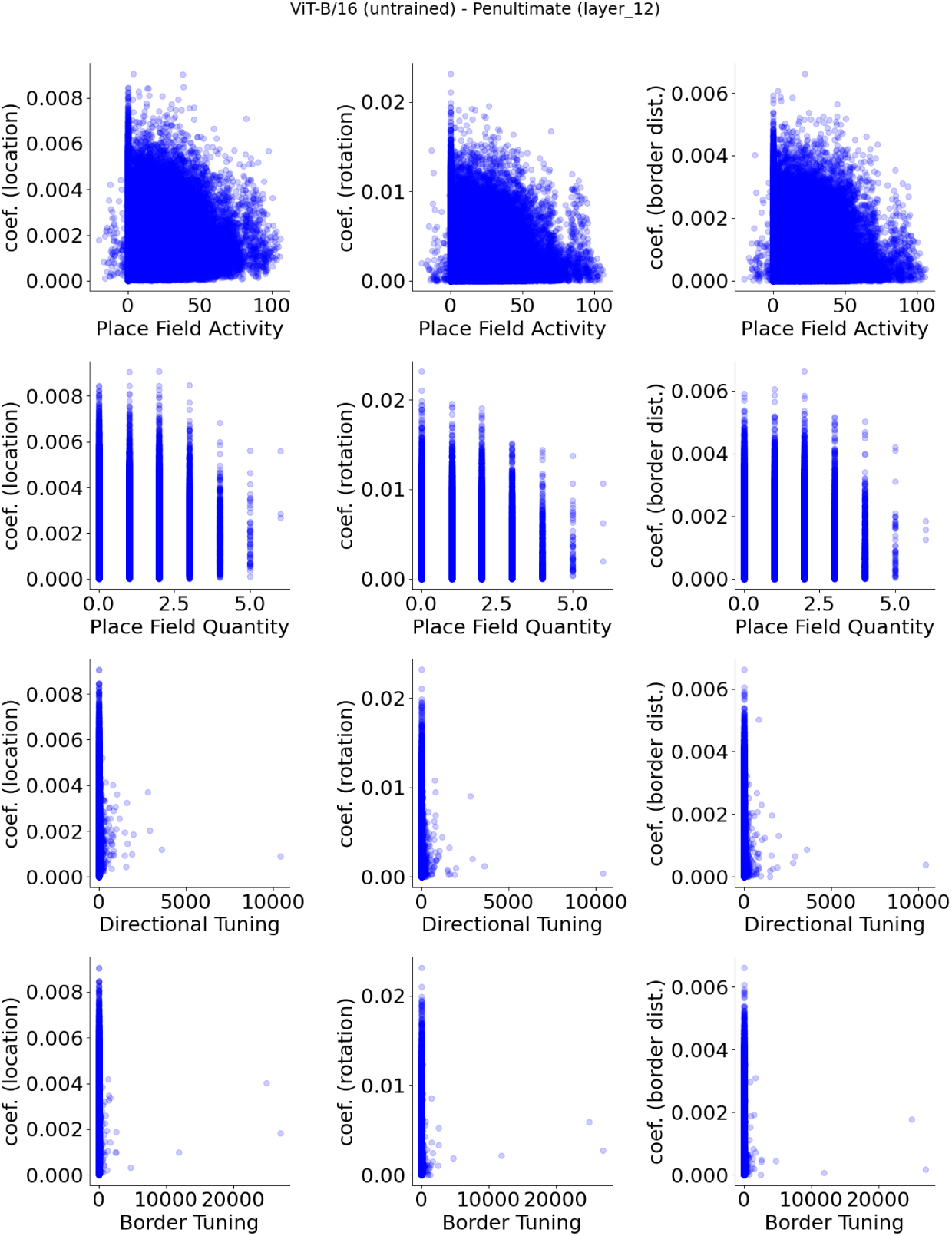
**ViT-B/16 (untrained), penultimate layer (layer 12) units, show no apparent relationship between spatial properties and their decoder weight strengths.**

**Figure S31:**
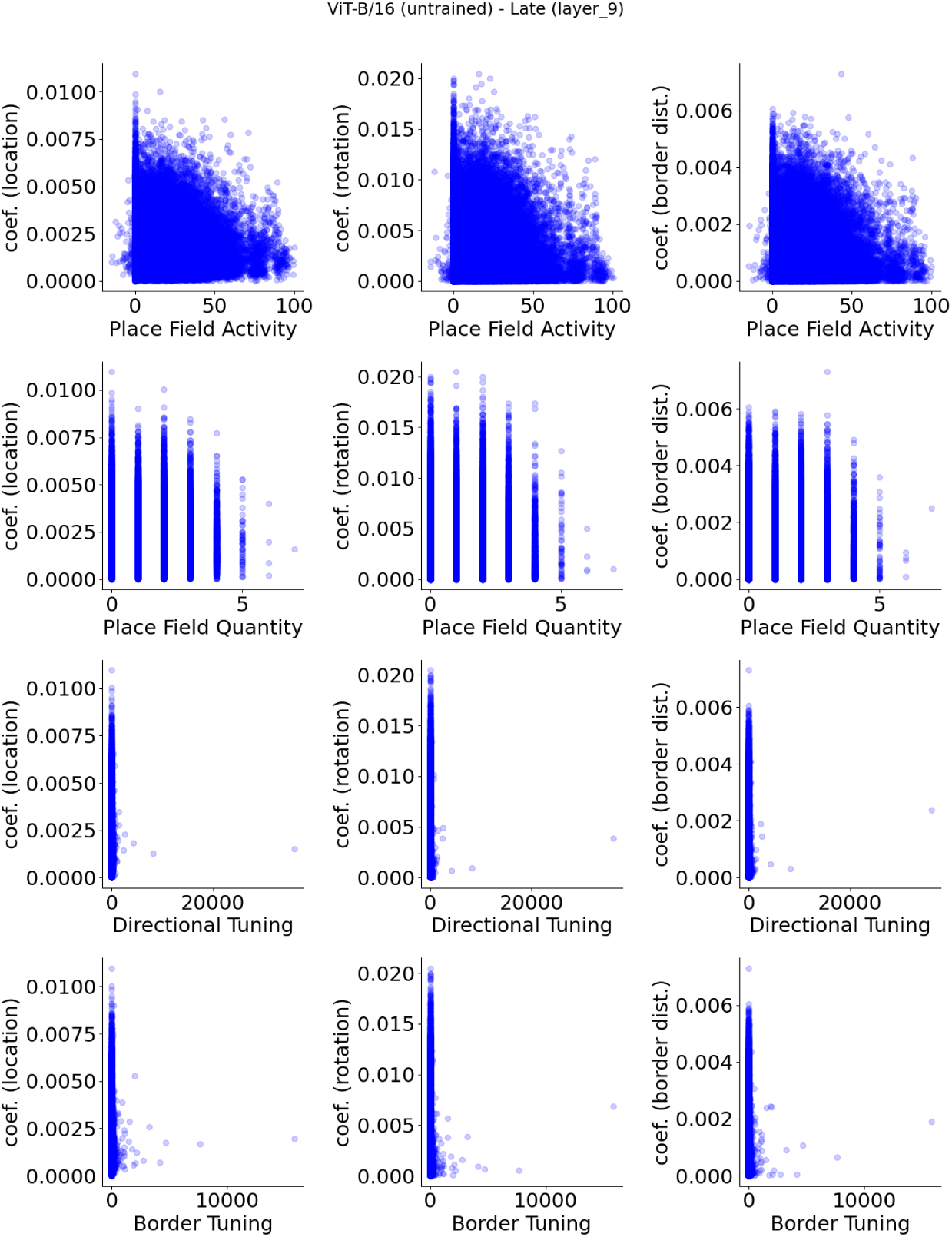
**ViT-B/16 (untrained), late layer (layer 9) units, show no apparent relationship between spatial properties and their decoder weight strengths.**

**Figure S32:**
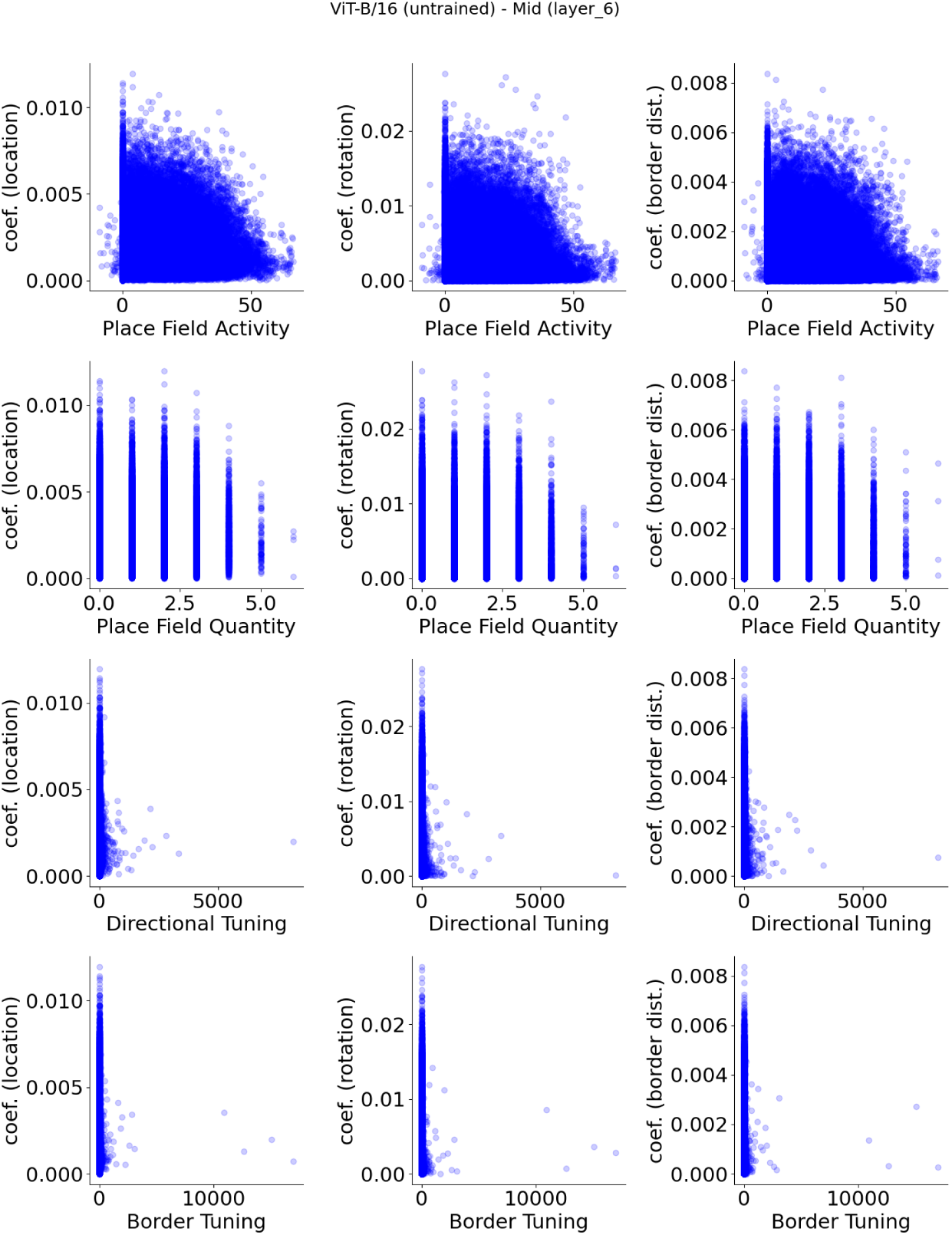
**ViT-B/16 (untrained), mid layer (layer 6) units, show no apparent relationship between spatial properties and their decoder weight strengths.**

**Figure S33:**
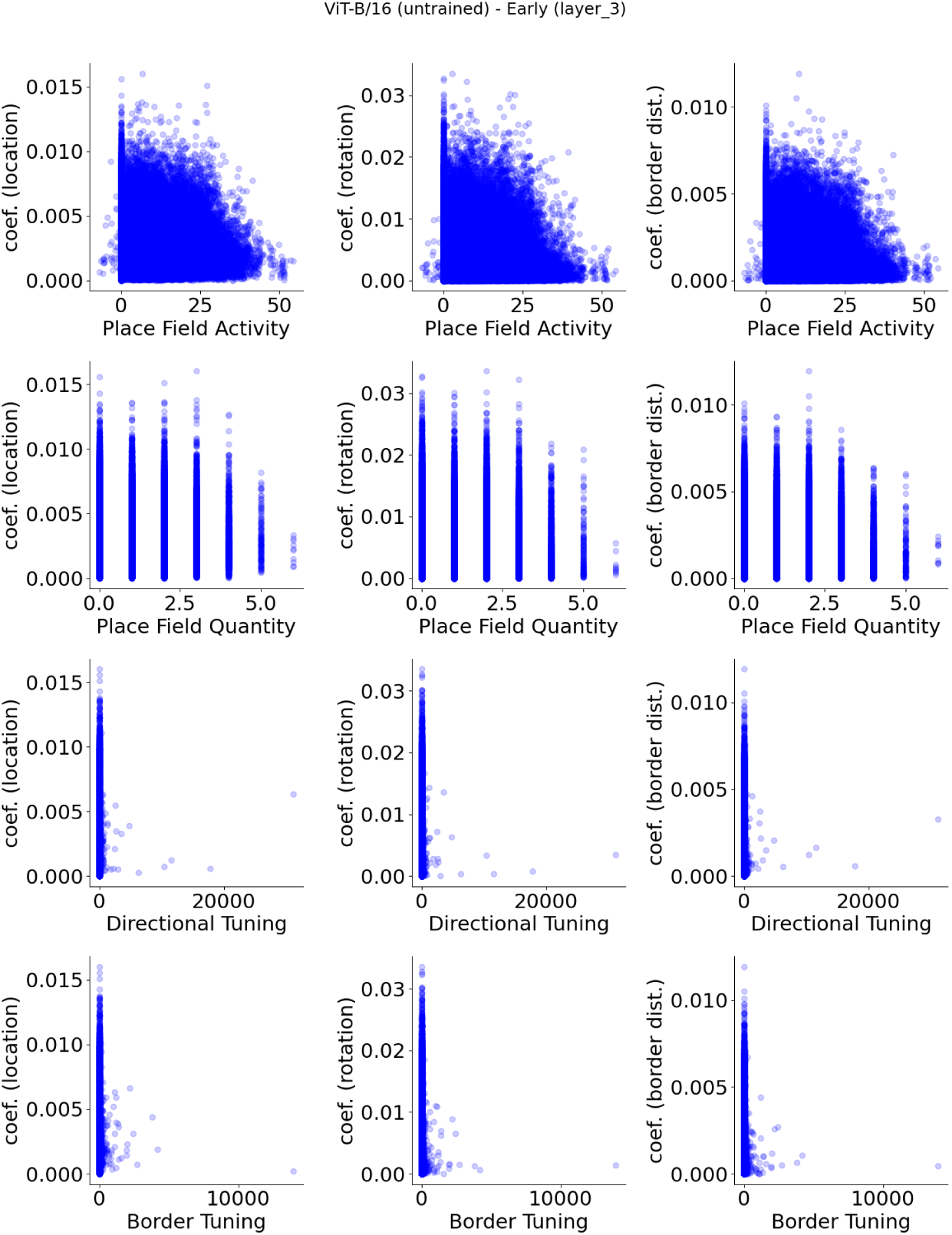
**ViT-B/16 (untrained), early layer (layer 3) units, show no apparent relationship between spatial properties and their decoder weight strengths.**

